# Cell type–resolved transcriptomic map of skeletal muscle in women with polycystic ovary syndrome

**DOI:** 10.64898/2026.03.13.711523

**Authors:** Tina Gorsek Sparovec, Gustaw Eriksson, Congru Li, Rutger Schutten, Haojiang Lu, Joana Rosa, Sara Torstensson, Sara Dahmani, Claes Ohlsson, Eva Lindgren, Pauliina Damdimopoulou, Angelica Lindén Hirschberg, Qiaolin Deng, Cecilia Lindskog, Elisabet Stener-Victorin

**Affiliations:** Department of Physiology and Pharmacology, Karolinska Institutet, Stockholm, Sweden; Department of Immunology, Genetics and Pathology, Uppsala University, Uppsala, Sweden; Department of Drug Treatment, Sahlgrenska University Hospital, Region Västra Götaland, Gothenburg, Sweden; Department of Internal Medicine and Clinical Nutrition, Institute of Medicine, Sahlgrenska Osteoporosis Centre, Centre for Bone and Arthritis Research at the Sahlgrenska Academy, University of Gothenburg, Gothenburg, Sweden; Department of Women’s and Children’s Health, Karolinska Institutet, Stockholm, Sweden; Department of Gynecology and Reproductive Medicine, Karolinska University Hospital, Stockholm

**Keywords:** PCOS, skeletal muscle, single nuclei RNA-seq, metformin, fibrosis

## Abstract

Polycystic ovary syndrome (PCOS) is associated with skeletal muscle insulin resistance, fibrosis, and lipotoxicity, yet the cellular origins remain unknown. Here, we present a comprehensive cellular atlas of skeletal muscle from hyperinsulinemic and hyperandrogenic women with PCOS and controls of similar age, weight, and BMI. Analysis of 72,247 nuclei from 19 biopsies revealed cell-type-specific dysregulation in PCOS, with fiber-type-specific metabolic impairment, converging with pro-fibrotic reprogramming of fibro-adipogenic progenitors (FAPs) and enhanced FAP-myofiber crosstalk, characterized by enhanced collagen and laminin signaling. Metformin intervention for 16-weeks selectively reversed PCOS-associated transcriptional dysregulation in FAPs, revealing heterogeneous cellular responses in skeletal muscle. *In vitro*, PCOS myotubes retained metabolic dysfunction, yet show normalized glucose responsiveness, indicating plasticity despite metabolic memory. Systemic hyperinsulinemia and hyperandrogenemia correlated with transcriptional signatures in muscle fibers and FAPs, linking endocrine imbalance to pro-fibrotic remodeling. These findings identify novel therapeutic targets beyond conventional insulin-sensitizing approaches.

**Highlights:** - First single-nuclei atlas of human skeletal muscle in women with PCOS

- FAPs subpopulations display activated pro-fibrotic phenotype

- Metformin selectively targets distinct cell populations

- Hyperinsulinemia and hyperandrogenemia correlate with transcriptional signatures of skeletal muscle fibers and FAPs

## Introduction

Polycystic ovary syndrome (PCOS), which affects approximately 15% of women worldwide, is a reproductive disorder characterized by hyperandrogenism and is closely linked to early-onset metabolic complications such as insulin resistance, type 2 diabetes, and cardiovascular disease.^1,2^ Hyperinsulinemia and tissue-specific insulin resistance occur in up to 90% of obese women with PCOS and 60% of lean women with PCOS.^1^ Unlike the global insulin resistance observed in type 2 diabetes, aberrant energy and glucose metabolism in PCOS is characterized by selective insulin resistance in skeletal muscle, adipose tissue, and liver, while ovarian and adrenal tissues remain insulin-sensitive.^3,4^ Skeletal muscle of women with PCOS exhibits impaired muscle glucose uptake, decreased mitochondrial oxidative phosphorylation (OXPHOS) activity, and downregulation of genes involved in OXPHOS. Indicating impaired mitochondrial function, as well as excessive production of reactive oxygen species (ROS), which further exacerbate insulin resistance.^5,6^ PCOS skeletal muscle shows markedly dysregulated gene expression involved in defective insulin signaling and tissue fibrosis, including TGF-β signaling, which may hinder insulin-stimulated glucose uptake and muscle remodeling, compounding insulin resistance.^7^ Additionally, skeletal muscle in PCOS shows altered global protein and DNA methylation expression pattern, as well as ectopic lipid accumulation leading to lipotoxicity.^6,8^ However, the specific cell types driving skeletal muscle pathology in PCOS, their molecular programs, and intercellular communication networks remain uncharacterized.

Treatment of PCOS is currently *ad hoc* and symptomatic, hindered by a lack of understanding of the underlying mechanisms. Lifestyle modification, mainly diet and increased physical activity, is the first line of treatment for women with PCOS and insulin resistance, followed by prescription of antidiabetic drugs such as metformin to regulate high blood glucose levels.^9,10^ While the exact mechanism of action still is debated, metformin is generally believed to indirectly activate AMPK by inhibiting OXPHOS complex I and thereby enhancing glucose and energy homeostasis within skeletal muscle.^11^ Besides energy-regulating effects, metformin has been demonstrated to attenuate fibrotic remodeling and improves mitochondrial function by targeting TGF-β signaling.^11,12,13,14^ Whether metformin has cell type specific effects in skeletal muscle is currently unknown.

In this study, we used single nuclei RNA sequencing (snRNA-seq) to generate the first transcriptional and spatial multiplex cellular atlas of skeletal muscle from overweight or obese, insulin-resistant and hyperandrogenic women with PCOS as well as from controls with regular menstrual cycle and similar age, weight, and BMI, and from women with PCOS again after a 16-week intervention with metformin. This approach enabled us to define cell type-specific molecular PCOS signatures and determine the restorative effect of metformin at the cellular level.

## Results

### Clinical characteristics

Biopsies of the vastus lateralis muscle of the quadriceps femoris were collected from 19 women with PCOS and 10 controls with similar age-, weight-, and BMI (Figure 1A, Table S1). After baseline examination, five women with PCOS were assigned to a 16-week intervention with metformin. At baseline, women with PCOS showed hyperandrogenism, with higher Ferriman-Gallwey scores (p < 0.001), androstenedione (p = 0.009), free androgen index (FAI) (p = 0.001), and lower SHBG (p < 0.001). In addition, they were hyperinsulinemic demonstrated by elevated fasting insulin (p = 0.003), fasting glucose (p = 0.05), homeostatic model assessment for insulin resistance (HOMA-IR) (p = 0.004), and area under the curve for insulin in the oral glucose tolerance test (p < 0.001). Women with PCOS also had increased circulating triglyceride levels (p = 0.024) and total cholesterol (p = 0.045). In the five women with PCOS, 16 weeks of metformin intervention did not result in significant hormonal changes.

**Figure 1:**
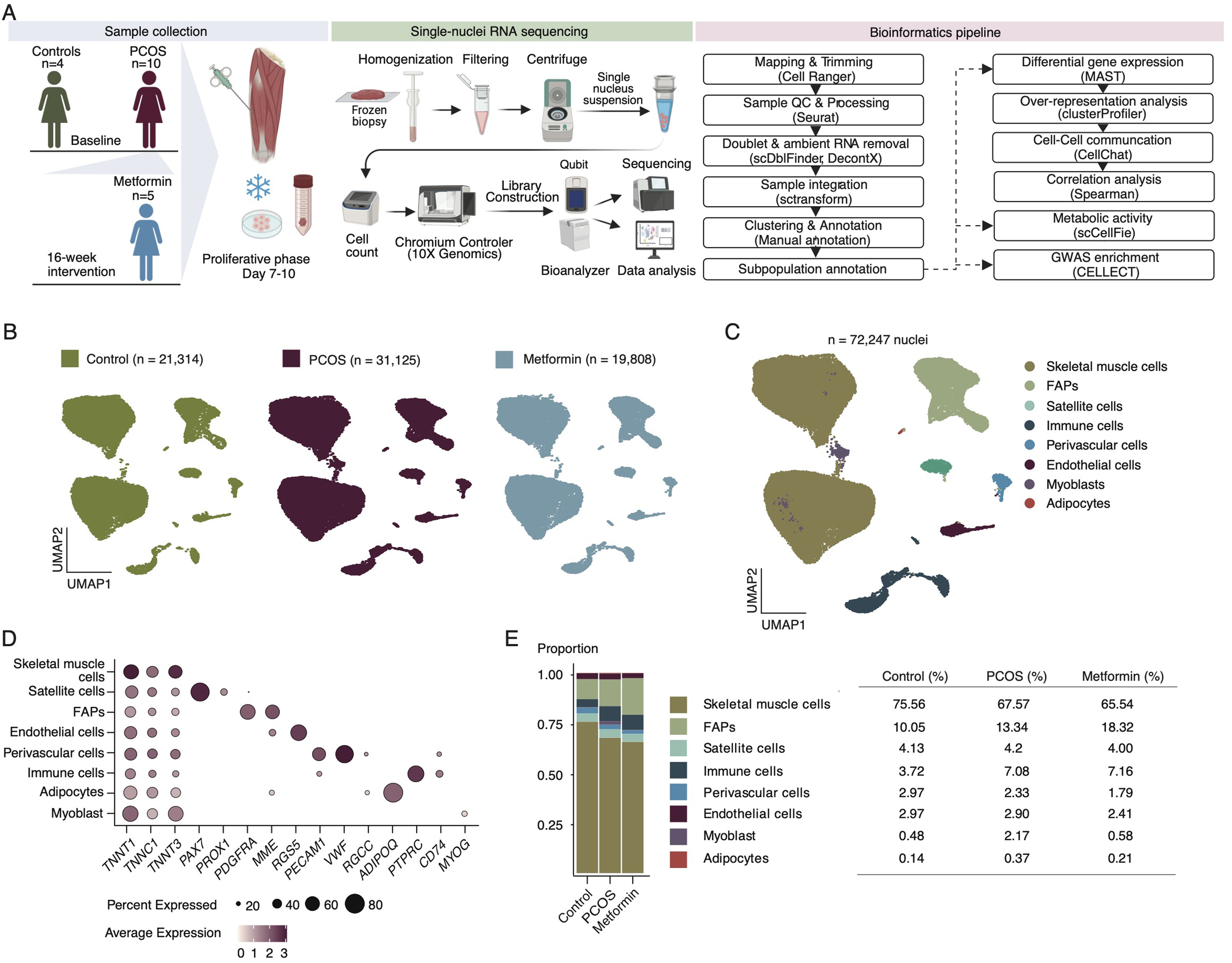
Single cell profiling of the skeletal muscle from women with and without PCOS. A) Illustration of the included participants, skeletal muscle biopsies, single nuclei RNA sequencing with 10X Genomics platform, bioinformatics analysis, and *in vitro* experiments using 2D skeletal muscle culture including Seahorse assay and bulk RNA sequencing using Prime-seq to validate the *in vivo* results. B) Distribution of nuclei in each group – Control, PCOS and PCOS after 16-weeks of treatment with metformin. C) Uniform manifold approximation and projection (UMAP) of integrated snRNA-seq data of 72,247 nuclei from 19 skeletal muscle samples revealed 8 major cell clusters. FAPs – fibro-adipogenic progenitors. D) Dotplot showing log-transformed gene expression of curated cell type markers. E) Barplot depicting the proportions of each major cluster across three groups; Control (n=4), PCOS (n=10), Metformin (n=5). E). Figure 1A created in BioRender.

### Transcriptomic landscape of the human PCOS-skeletal muscle

We generated the first single-nuclei transcriptomic atlas of human skeletal muscle biopsies from women with PCOS (n = 10) and controls (n = 4) to identify cell type-specific disease signatures, and to define the effect of 16-week metformin treatment on transcriptomic reversibility in women with PCOS (n = 5) using the 10X Genomics protocol (Figure 1A). After quality control, stringent filtering and ambient RNA removal, 72,247 nuclei were captured from the 19 sequenced samples: Control = 21,314, PCOS = 31,125, and Metformin = 19,808 (Figures 1B, S1A-C Table S2). Eight main clusters were identified based on the markers previously used in the literature^15–17^ (Figures 1C-D, S1D): skeletal muscle cells (n = 50,117), fibro-adipogenic progenitors (FAPs) (n = 9,921), satellite cells (n = 2,994), immune cells (n = 4,414), perivascular cells (n = 1,712), endothelial cells (n = 2,013), myoblasts (n = 890) and adipocytes (n = 186) (Table S3). No differences were detected in the relative abundance of nuclei in the main clusters among the three groups, indicating cell proportion stability across conditions (Figures 1D-E, S1E Table S4).

### PCOS skeletal muscle cells exhibit impaired metabolic flexibility

Subcluster analysis of skeletal muscle cells identified four subpopulations based on the mentioned marker genes: MF-I (muscle fiber type I), MF-IIa (muscle fiber type IIa), MF-IIx (muscle fiber type IIx), and Hybrid I-II muscle fibers, the latter expressing genes from both type I and II fibers (Figures 2A-B, S2A, S1H).^15–18^ There were no differences in the proportion of nuclei between skeletal muscle cell subpopulations in PCOS compared with control (Figures 2C, S2B).

**Figure 2:**
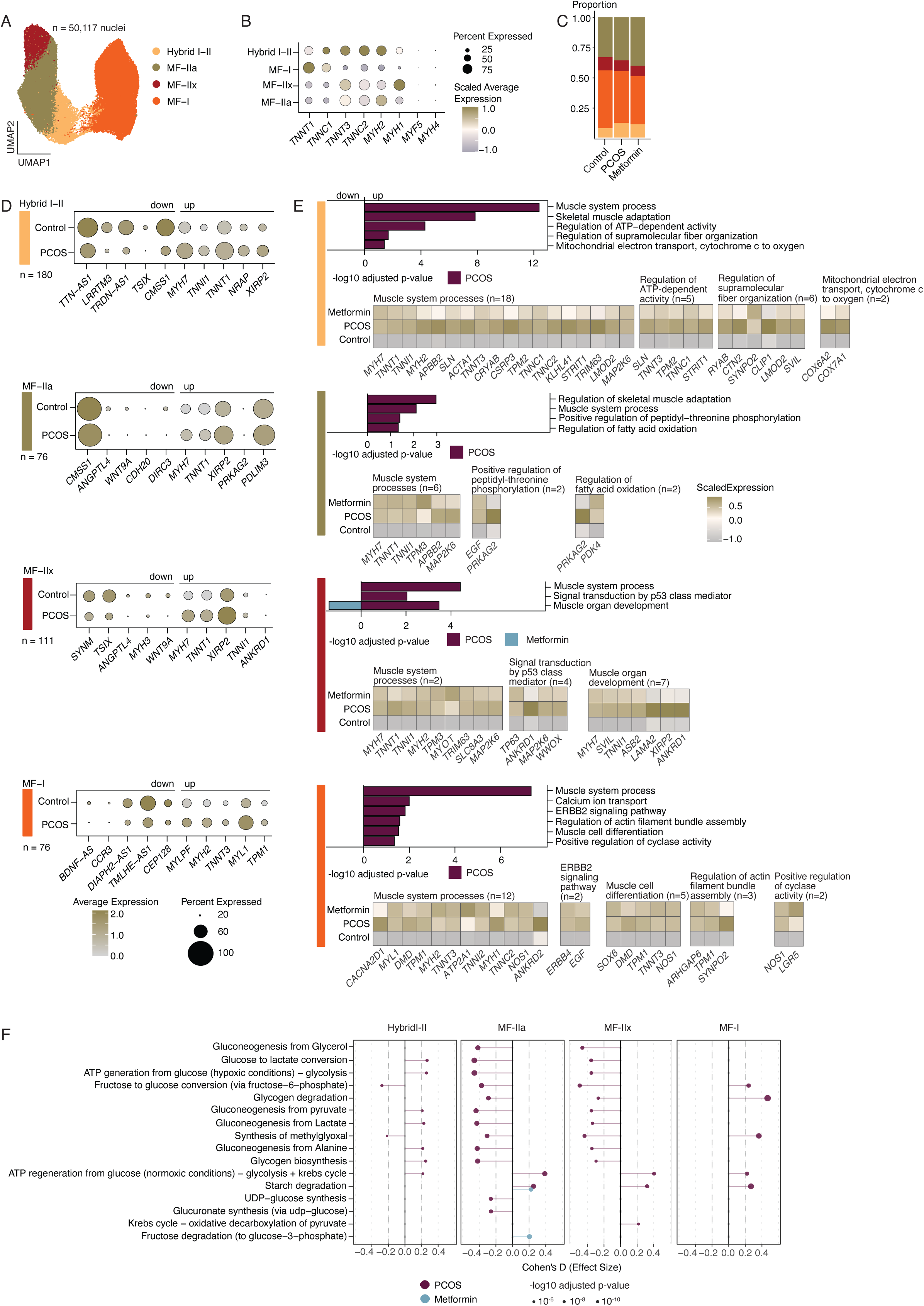
Transcriptomic characterization of skeletal muscle cluster and the effect of 16-week metformin treatment in women with PCOS. A) Uniform manifold approximation and projection (UMAP) of 50,117 nuclei from all patients revealed four subclusters: MF-I (muscle fiber type I), MF-IIx (muscle fiber type IIx), MF-IIa (muscle fiber type IIa), and Hybrid I-II muscle fibers. B) Dotplot showing log-transformed gene expression of skeletal fiber marker genes across identified muscle fiber types. C) Barplot depicting the proportions of the four skeletal muscle fiber type in Control, PCOS and Metformin group. D) Top 5 of up-and downregulated DEGs (Control vs. PCOS) with the total number of DEGs identified in each skeletal muscle fiber type. E) Gene Ontology (GO) enrichment analysis on DEGs within each subcluster at baseline (Control vs. PCOS) and after intervention (Metformin vs. PCOS). Heatmaps under the GO show the DEGs in corresponding GO terms, presenting the scaled expression in Control, PCOS and Metformin groups. Significant GO terms with Benjamini-Hochberg adjusted p-value < 0.05 are visualized with -log_10_ transformed values. F) Differential metabolic tasks in skeletal muscle fibers comparing PCOS to Control and Metformin to PCOS. Lollipop plot shows Cohen’s d effect size for glucose and glycogen metabolism tasks. Dot size indicates statistical significance (-log_10_ adjusted p-value. Dashed vertical lines mark effect size threshold + 0.2. Negative values: downregulated in PCOS, Positive values: upregulated in PCOS.

Differentially expressed gene (DEG) analyses of each subpopulation comparing women with and without PCOS identified most DEGs in the Hybrid I-II muscle fiber type (n = 180), followed by MF-IIx (n = 111), MF-IIa (n = 76), and MF-I (n = 76) subclusters (Figure 2D, Tables S5-6). Several of the top five up- and downregulated DEGs, such as *XIRP2*, *WNT9A*, and *ANGPTL4* (Figure 2D), were shared across muscle fiber subpopulations, suggesting a common effect of PCOS on the skeletal muscle fiber transcriptome. We observed a decreased expression of *ANGPTL4*, a member of the angiogenin-like protein family, in MF-IIx and MF-IIa. *ANGPTL4* expression has previously been shown to positively correlate with body fat and insulin resistance and negatively correlated with lean mass in skeletal muscle.^19^ MF-IIx and MF-IIa fibers exhibit significantly reduced expression of *WNT9A*, a gene suggested to play a role in muscle cell differentiation *in vitro* and reported to be upregulated in skeletal muscle following sprint exercise (Figure 2D).^20,21^ *XIRP2*, a gene encoding Xin actin-binding repeat containing 2, which stabilizes the F-actin structure, and ensures structural integrity,^22^ is upregulated in MF-IIx, MF-IIa, and Hybrid I-II fibers, whereas it was downregulated in previous bulk transcriptomic and proteomic analyses of PCOS skeletal muscle.^8^ Additionally, canonical marker genes for defined fiber subtypes including *TNNT1*, *MYH2*, *TNNT3*, and *TNNI1,* were differentially expressed suggesting altered sarcomere gene programs in PCOS fiber subpopulations. Mutations in these sarcomere genes have been associated with hypertrophic cardiomyopathy risk, which is elevated in PCOS.^23,24^ In MF-I fibers, downregulation of *BDNF-AS* (Brain-Derived Neurotrophic Factor Antisense RNA), a negative regulator of BDNF expression, together with upregulation of *TPM1,* may increase BDNF bioavailability and alter contractile filament regulation, potentially affecting muscle plasticity and muscle contraction, respectively.^25,26^

Gene Ontology (GO) analysis of the DEGs in each skeletal muscle cell subtype revealed significant enrichment of the biological processes (BP) related to upregulated pathways in skeletal muscle system process, muscle organ development, and muscle cell differentiation in PCOS compared to controls (Figures 2E, S2E, Tables S7,8). These transcriptional alterations are consistent with the effect of hyperandrogenism, a characteristic feature of PCOS, which has been linked to increased skeletal muscle mass.^27^ We also observed upregulation of enriched pathways related to metabolism, including changes in mitochondria and fatty acid oxidation in the Hybrid I-II and MF-IIa subpopulations, respectively (Figure 2E). Among the upregulated enriched genes in Hybrid I-II, we identified cytochrome c oxidases *COX7A1* and *COX6A2* which are typically induced under hypoxic conditions via HIF1-a, suggesting hypoxia-dependent reprogramming of PCOS Hybrid I-II.^28^ Such reprogramming promotes can elevate oxidative stress through increased ROS production, which has been linked to impaired insulin signaling in PCOS skeletal muscle.^4,5,28,29^ Upregulated protein kinase AMP-activated non-catalytic subunit gamma 2 (*PRKAG2)* and pyruvate dehydrogenase kinase 4 (*PDK4)* in MF-IIa indicate suppressed glucose oxidation and preferential fatty acid utilization, consistent with metabolic inflexibility observed in insulin-resistant skeletal muscle in mice.^30^ Genome-wide association studies (GWAS) have demonstrated that PCOS is associated with alterations in the ERBB2 signaling pathway, a regulator of cell proliferation implicated in granulosa cell dysfunction and hormonal imbalances.^31^ However, in muscle, ERBB2 is crucial for muscle regeneration and muscle spindle maintenance.^32^ Metformin reversed enriched BP linked to muscle organ development only in MF-IIx, indicating cell-type-specific action of metformin. Whereas in other subpopulations it showed reversal of individual genes only including *CTN2, CLIP1, LAMA2, PRKAG2*, *ANKRD2*, and *CACNA2D1* without pathway-level normalization (Figure 2E). This heterogenous pattern suggests selective rather than uniform metformin sensitivity across muscle fiber subtypes. Among the normalized genes *ANKRD2* has been shown to decrease with metformin treatment in rat muscle arteries, and *PRKAG2* polymorphisms have been identified as predictors of metformin response.^33,34^ These findings demonstrate fiber-type-specific transcriptional dysregulation in PCOS, with metformin inducing targeted gene-level changes rather than comprehensive pathway normalization.

To further investigate the metabolic landscape in PCOS skeletal muscle we used scCellFie which infer metabolic task activity based on gene-protein-reaction rules.^35^ We observed significant dysregulation in metabolic tasks in MF-IIa (n = 71 metabolic tasks), Hybrid I-II (n = 50), MF-IIx (n = 41), and MF-I (n = 25) in PCOS compared with controls (Figure S2C). PCOS MF-IIa and MF-IIx fibers showed downregulated ATP production from glycolysis, gluconeogenesis, and glycogen biosynthesis, while ATP production via oxidative pathway, and starch degradation were upregulated compared with controls (Figure 2F), suggesting impaired glucose handling and a shift in fuel utilization. In MF-I fibers, scCellFie inferred upregulated glycogen degradation and methylglyoxal synthesis, which is responsible for the formation of advanced glycation end-products and is recognised as a causal mediator of insulin resistance,^36^ along with decreased ROS detoxification. This metabolic signature is consistent with elevated oxidative stress which has been previously reported in PCOS skeletal muscle.^3^ In MF-IIa fibers, metabolic tasks related to amino acid turnover and fatty acid synthesis were downregulated (Tables S9-S10). Reduced fatty acid synthesis, coupled with the upregulation of *PDK4* and *PRKAG2* described above, suggest a coordinated switch towards fatty acid oxidation at the expense of glucose utilization. We also observed strong suppression of glutamine synthesis, potentially limiting glutamine-dependant processes. Notably, glutamine is also required for collagen synthesis which we observed altered in FAPs (see below). In Hybrid I-II fibers we observed small-to-moderate upregulation of sphingolipid and ceramide synthesis (Tables S9-S10), which have been previously linked to lipid-induced insulin resistance, further confirming fiber-type-specific metabolic dysregulations and impaired insulin sensitivity.^37,38^ Metformin treatment did not induce profound changes toward improving dysregulated metabolic tasks at the transcriptional level, suggesting limited direct metformin action on PCOS skeletal muscle, although mechanism such as AMPK activation, which are not captured by transcriptomics, may still contribute to the metabolic improvements.

### *In-vitro* validation of PCOS skeletal muscle dysfunction

To investigate the impact of PCOS on skeletal muscle cell function and to validate the findings from snRNAseq, we cultured isolated satellite cells in differentiation medium with and without metformin for 8 days before performing prime-seq, Seahorse assay, glucose uptake assay and immunofluorescence staining (Figure 3A).

**Figure 3:**
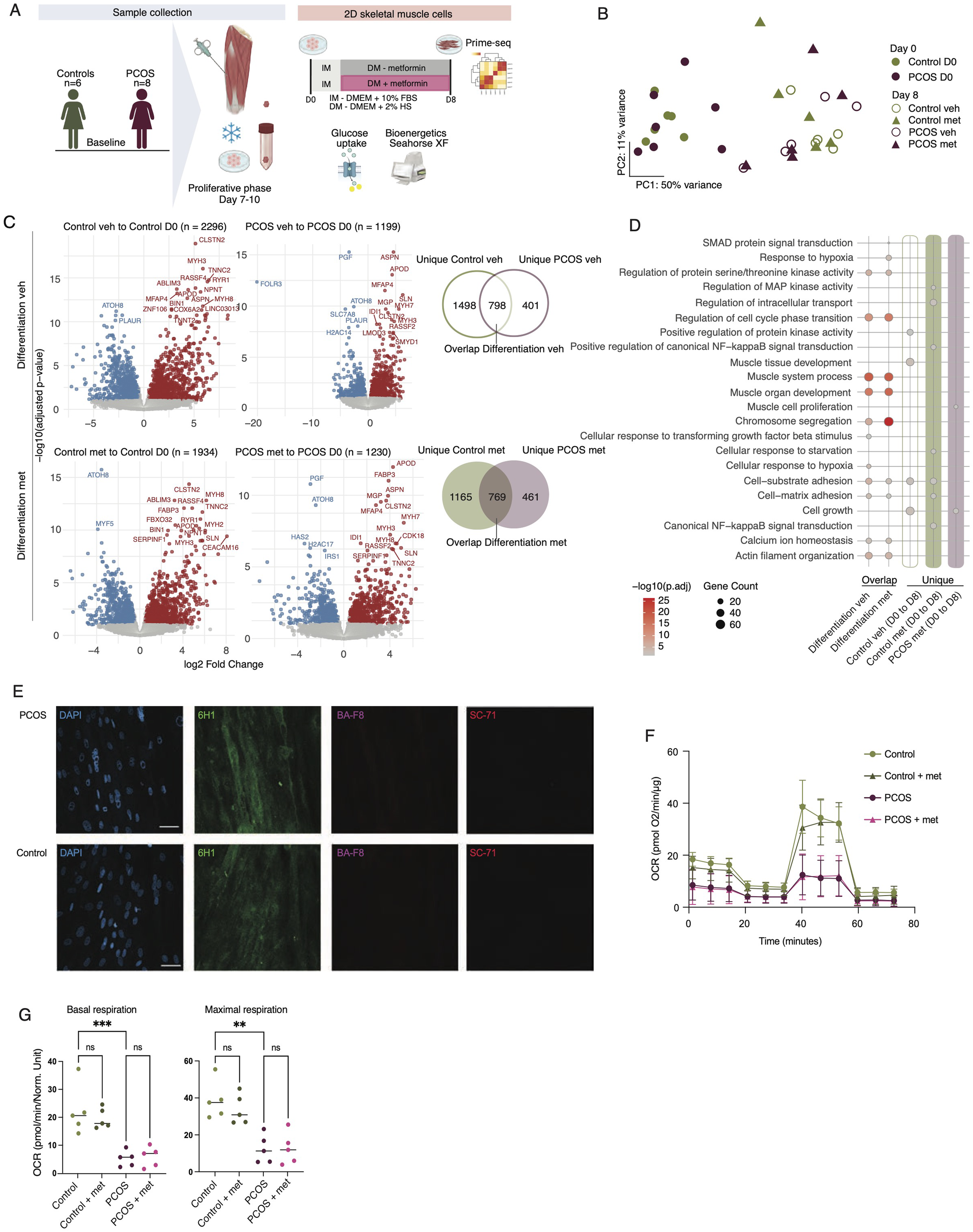
*In vitro* validation using 2D skeletal muscle cells. A) Illustration of the experiment design and 2D skeletal muscle cell culturing conditions B) Principal component analysis (PCA) of variance-stabilized gene expression demonstrates clear separation between D0 and D8 samples, without clear separation of PCOS and Controls treated with (met) or without (veh) metformin. C) Vulcano plots illustrate differential gene expression between undifferentiated (D0) and differentiated (veh/met) conditions, with upregulated and downregulated genes log_2_ Fold Change (X-axis) and -log_10_ adjusted p-value (Y-axis). Venn Diagrams represent shared and condition-specific transcriptional signature -unique for both vehicle and metformin treated cells. D) Selected GO terms for the corresponding enrichment of shared and unique DEGs in both vehicle and metformin treated cells. E) Representative immunofluorescence staining of fixed differentiated skeletal muscle fibers from one PCOS and one control sample stained with 6H1 (Type IIx), BA-F8 (Type I), and SC-71 (Type IIa), to identify muscle fiber type composition. DAPI stained nuclei. Scale bars are 50 μm. F) Oxygen consumption rate (OCR) measured in differentiated PCOS and Control myotubes with and without metformin (n=5/group). Data are presented as mean ± SD. G) Basal and maximal respiration measured in OCR. Each dot represents a sample; bar is a mean. \*\**p* < 0.01, \*\**p* < 0.001 using ordinary one-way ANOVA with Tukey correction for multiple comparisons.

#### Transcriptional signature of cultured myotubes

Principal component analysis showed tight clustering at Day 0 and Day 8, regardless of treatment or sample group (Figure 3B). There were no DEGs between PCOS and controls at Day 0, while at Day 8, *FOLR3, RPL10P6* and *RPL10P9* were differentially expressed (Figure S2D). Treatment with 1 mM metformin led to upregulation of *POU2F2* and *RPL10P9* in PCOS cells. In the control group, metformin exposure affected gene expression related to oxidative phosphorylation, and steroid hormones (n = 12 DEGs, Table S11). In contrast, myotubes from PCOS treated with metformin significantly upregulated *FOLR3* and downregulated *TXNIP* (Table S10). Since no DEGs were detected at Day 0, the analysis focused on differences in differentiation. In the vehicle group, we identified 798 overlapping genes between PCOS and controls at Day 8 (Figure 3C), predominantly associated with biological processes involving cell cycle regulation, muscle system processes, chromosome segregation, cell-substrate adhesion and calcium ion homeostasis (Figure 3D, Table S12). Enrichment analysis of 1498 unique DEGs in the control group without metformin treatment indicated differences in skeletal muscle development, cell growth, and regulation of protein kinase activity (Table S13). By contrast, no significant enrichment was observed in the PCOS group, suggesting aberrant capacity for muscle growth and decreased protein kinase activity *in vitro*. In the metformin group, 769 overlapping DEGs were enriched for muscle system processes, chromosome segregation, SMAD signal transduction, cell substrate adhesion, response to hypoxia and calcium ion homeostasis (Figures 3C, D, Table S12). The 1165 unique DEGs in the control metformin group were enriched for regulation of MAP kinase, NF-Kb signal transduction and cell-substrate adhesion, pathways previously discovered to be targeted by metformin.^39^ 461 unique DEGs in the PCOS metformin group were enriched for cell growth and muscle cell proliferation (Figure 3D, Table S13). This aligns with our snRNA-seq finding showing metformin-induced reversal of muscle organ development processes specifically in MF-IIx fibers, suggesting that metformin engages in proliferative and repair-related programs in PCOS muscle. Overlapping DEGs and corresponding biological processes in both vehicle and metformin groups show a consistent muscle development signature, with minimal impact of metformin.

#### Differentiated myotubes display type MF-IIx phenotype

To confirm successful differentiation in myotubes, we examined the expression of established skeletal muscle fiber and myogenic markers. Differentiated cells in both vehicle and metformin group show upregulation of both slow and fast twitch fiber related genes, with no differences between PCOS and control (Figure S2E). These transcriptomics findings were further validated at the protein level, revealing a MF-IIx phenotype in both PCOS and control myotubes (Figure 3E).

#### Myotubes display impaired oxidative metabolism but preserved glucose sensitivity

Mitochondrial oxygen consumption was assessed using the Seahorse XFe96 Analyzer on differentiated myotubes with and without metformin treatment. Basal respiration was significantly lower in PCOS myotubes (5.2 ± 2.8 pmol/min/µg) compared to control myotubes (22.3 ± 8.9 pmol/min/µg; p < 0,006, Figures 3E-F). Similarly, maximal respiration was lower in PCOS myotubes (12.4 ± 7.6 pmol/min/µg) compared to controls (36.6 ± 10.2 pmol/min/µg, p < 0,001; Figures 3F, G), indicating impaired mitochondrial capacity. While we observed retained mitochondrial phenotype in PCOS myotubes *in vitro*, the glucose uptake assay revealed no differences in glucose uptake between PCOS and control myotubes with and without insulin (Figure S2). A 1 mM metformin treatment during differentiation did not have any significant effect on OCR in either control (control + met) or PCOS (PCOS + met) myotubes (p > 0.05) (Figure 3F-G), suggesting that prolonged metformin treatment did not improve mitochondrial function in either condition.

### Pro-fibrotic dysregulations in FAP subtypes are effectively reversed by metformin

Skeletal muscle FAPs constitute a heterogeneous population, exhibiting distinct phenotypes and functional roles *in vivo*.^40^ This diversity is influenced by dynamic changes in the microenvironment and is further modulated under pathological conditions.^41^ Here, we identified three FAP subpopulations: GPC3^+^, MME^+^, and FBN1^+^ FAP, each linked to specific functions (Figures 4A-B, S4A).^16,42,43^ Pro-adipogenic MME^+^ FAPs strongly express *PDGFRA*, *MME* and *DCN*. In contrast GPC3+ FAPs display a distinct signature defined by *GPC3, GLI1* and *FBLN1*, and are associated with reduced adipogenic potential. FBN1^+^ FAPs express canonical markers *FBN1*, and *CD55*, suggesting roles in extracellular matrix (ECM) organization. Previously reported markers *LUM*, and *THY1* show low expression in the FAPs in our data set (Figure S1H).

**Figure 4:**
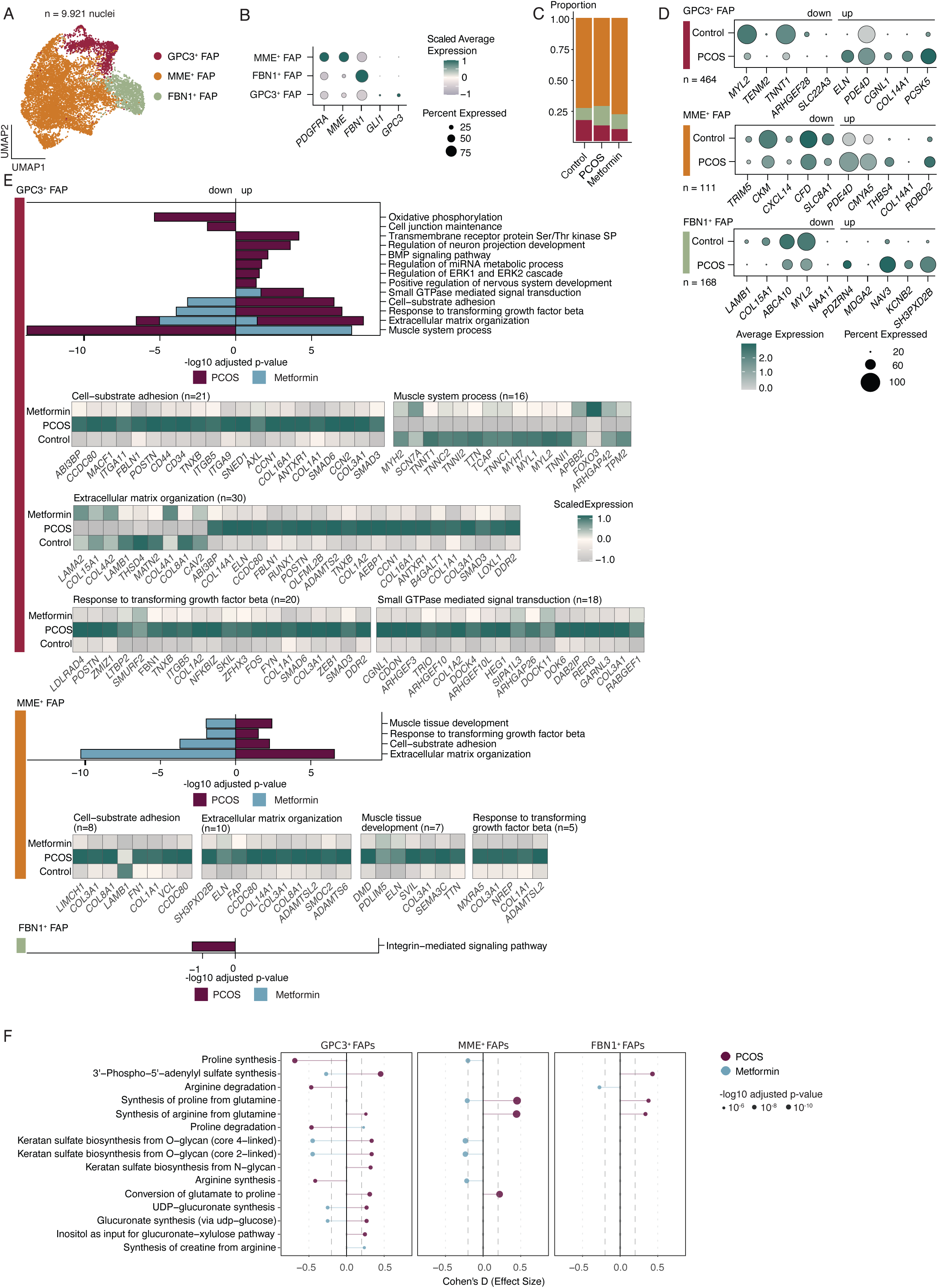
Transcriptomic characterization of fibro-adipogenic progenitor cluster and the effect of 16-week metformin treatment in women with PCOS. A) Uniform Manifold Approximation and Projection (UMAP) of 9,921 nuclei from all patients revealed three distinct FAP subclusters: MME^+^, FBN1^+^ and GPC3^+^ FAP. B) Dotplot showing log-transformed gene expression of FAP marker genes across identified FAPs types. C) Barplot depicting the proportions of the FAP subclusters in Control, PCOS and Metformin group. D) Top 5 of up-and downregulated DEGs (PCOS vs. Control) presented in dotplot in average expression with the total number of DEGs identified in each FAP subcluster. E) Gene Ontology (GO) enrichment analysis on DEGs within each subcluster at baseline comparing PCOS and Control (PCOS); and after 16-week intervention comparing Metformin vs. PCOS (Metformin). GOs with Benjamini-Hochberg adjusted p-value < 0.05 are reported and -log_10_ transformed for visualization. Heatmaps under the GO show the DEGs in corresponding metformin-reversed GO terms. F) Differential metabolic tasks in FAPs comparing PCOS to Control and Metformin to PCOS. Lollipop plot shows Cohen’s d effect size for metabolism tasks related to proteoglycan synthesis and glycosylation. Dot size indicates statistical significance (-log10 adjusted p-value). Dashed vertical lines mark effect size threshold ± 0.2. Negative values: downregulated in PCOS, Positive values: upregulated in PCOS.

Analysis of cluster composition revealed no differences in proportions across the groups (Figures 4C, S4B). To investigate transcriptional changes associated with PCOS, we performed DEG analyses in identified FAP subpopulations. GPC3^+^ FAPs displayed the most DEGs (n = 464) followed by FBN1^+^ (n = 168) and MME^+^ FAPs (n = 111) (Figure 4D, Tables S5-S6). In GPC3^+^ FAPs, downregulation of muscle associated genes such as *ARHGEF28*, *TNNT1*, *MYL2* suggests a compromised support of muscle function and integrity.^44^ In contrast, upregulation of *ELN*, *PDE4D*, *COL14A1* and *PCSK5* reflects an activated, pro-fibrogenic state of GPC3^+^ FAPs which in previous studies has been demonstrated to be contributing to the pathological muscle remodeling.^45–47^ In MME^+^ FAPs we detected downregulation of *CKM*, encoding muscle-specific creatine kinase, suggesting a decreased ATP synthesis capacity and thus potentially compromised energy metabolism in PCOS FAPs.^48^ Downregulation of several immune-related genes such as *TRIM5*, *CXCL14,* (a chemokine involved in macrophage recruitment) and *CFD* (complement factor d, a component of the complement system that also regulates adipogenesis),^49,50,51^ suggesting potential alterations in MME+ FAPs-immune cell crosstalk in PCOS (Figure 4D). Decreased expression of *SLC8A1*, a sodium/calcium exchanger, indicates a decreased capacity for intracellular calcium dynamics and could contribute to insulin resistance.^52^ Like in GPC3^+^ FAPs, we detected an upregulation of ECM components *PDE4D*, *CMYA5*, *THBS4*, and *COL14A1* as well as *ROBO2*, a gene encoding a receptor involved in cell migration, which all together might impact ECM organization and stability, and impaired cell migration.^53,54^ Similarly, the top 10 DEGs in FBN1^+^ FAPs showed downregulated *LAMB1*, and *COL15A1*, and upregulation of *MDGA2*, *NAV3* and *SH3PXD2B* which are implicated in regulating cell adhesion, and cell mobility.^55,56^ COL15A1 has also been reported downregulated in fibrotic models, suggesting disruption in structural integrity.^57^ Downregulation of *ABCA10*, might impact lipid homeostasis. Dysregulation of *MYL2*, *PDZRN4* and *KCNEB2* has been shown to influence contractile function, as well as protein turnover and membrane potential.^58^ Together, these dysregulations indicate a contribution of FBN1^+^ FAPs to muscle structure impairment.

In GPC3^+^ FAPs, GO term analysis for BP of DEGs between PCOS and controls revealed dysregulations in biological processes linked to muscle tissue function, ECM organization, TGF-β response, and cell-substrate adhesion, which were normalized by metformin (Figure 4E, Tables S7-S8). A similar pattern was observed in pro-adipogenic MME^+^ FAPs, suggesting that PCOS milieu induced a switch toward a more fibrotic phenotype in both GPC3^+^ FAPs and MME^+^ FAPs with metformin exerting a restorative effect on key structural and signaling pathways involved in fibrosis, as previously suggested in cardiac fibrosis.^59^ In contrast, in GPC3^+^ FAPs, perturbations in key canonical metabolic components: oxidative phosphorylation, BMP signaling, Ser/Thr kinase, and ERK cascade were not fully reversed to the control levels by metformin. However, certain genes like *SMAD6, CNN1* (BMP signaling pathway), *CNN1/2,* and *DAB2IP* (regulation of ERK1/ERK2 cascade) and *ZEB1* (Ser/Thr kinase signaling pathway) were still restored by metformin (Figure S4C). This suggest that in GPC3^+^ FAPs metformin targets both canonical and noncanonical TGF-β signaling pathways and their downstream transcriptional programs.^60^ Distinct from GPC3^+^ and MME+ FAPs, FBN1^+^ FAPs, which are responsible for ECM organization, displayed a downregulated integrin-mediated signaling pathway in PCOS, suggesting potential alterations in adhesion, or differentiation capacity (Figure 4E).

Next, inferred metabolic task activity in FAPs exhibited a profound metabolic heterogeneity across FAP subpopulations, with GPC3^+^ FAPs as the most dysregulated (n = 100 metabolic tasks), followed by MME^+^ (n = 42) and FBN1+ FAPs (n = 39) (Figure S4D). Predominantly in GPC3^+^ and MME^+^ FAPs, a metabolic signature centers on glutamine metabolism. Both glutamate and glutamine synthesis and degradation are markedly suppressed (Cohen’s d < - 0.5) in GPC3^+^ FAPs, suggesting a metabolic channelling of glutamine and glutamate to ECM precursors, rather than active turnover. Multiple tasks related to glutamine conversion to collagen precursors are upregulated, such as synthesis of proline, ornithine, and arginine with a larger effect size in MME^+^ FAPs compared to GPC3^+^ and FBN1^+^ FAPs (Figures, 4D, S4E, Tables S9b). Additionally, hexosamine biosynthetic tasks, including N-acetylglucosamine (GlcNAc) and UDP-N-acetyl-D-galactosamine (UDP-GalNAc) synthesis, was strongly upregulated in both MME^+^ and GPC3^+^ FAPs, providing essential substrate for the downstream N-linked glycosylation required for ECM production. These results are consistent with previous report of reduced glutamine levels and increased UDP-GlcNAc levels in obese adipocytes.^61^ Furthermore, GPC3^+^ FAPs display a robust upregulation of the ECM biosynthetic machinery with small-to-medium effect size for proteoglycan production via upregulated keratan sulfate biosynthesis tasks; and for N-linked glycosylation, which is essential for proper folding and secretion of ECM proteins (Figure 4F). Metformin treatment effectively restored proteoglycan biosynthesis and glycosylation tasks, whereas glutamine synthesis and conversion to collagen precursors remained largely unaffected (Figure 4F, Table S10). Collectively, these data show that transcriptional remodeling in FAPs is subpopulation-specific, with a shift toward a pro-fibrotic phenotype and dysregulated glutamine metabolism in MME^+^ and GPC3^+^ FAPs, representing the most pronounced disease-associated alterations.

### Dysregulated PAX7⁺ and ENO3⁺ satellite subpopulation in PCOS muscle

Recent studies have shown heterogeneity among satellite cells in skeletal muscle based on their maturation stage.^15,62^ In our dataset of 2,994 satellite cell nuclei, we identified three distinct clusters: PAX7^+^ MSC, which strongly express *PAX7*, *IGFBP5*, and *NCAM1* but have low *MYF5* expression; PAX7^-^ MSC, with low *PAX7* expression; and ENO3^+^ MSC, which express typical satellite cell genes such as *PAX7*, along with higher expression of skeletal muscle-specific markers such as *TNNC3* and *MYH2,* indicating activated myogenic program (Figures 5A-B, S1H).^15^ Analysis of cluster composition revealed no differences in proportions across the groups (Figure 5C).

**Figure 5:**
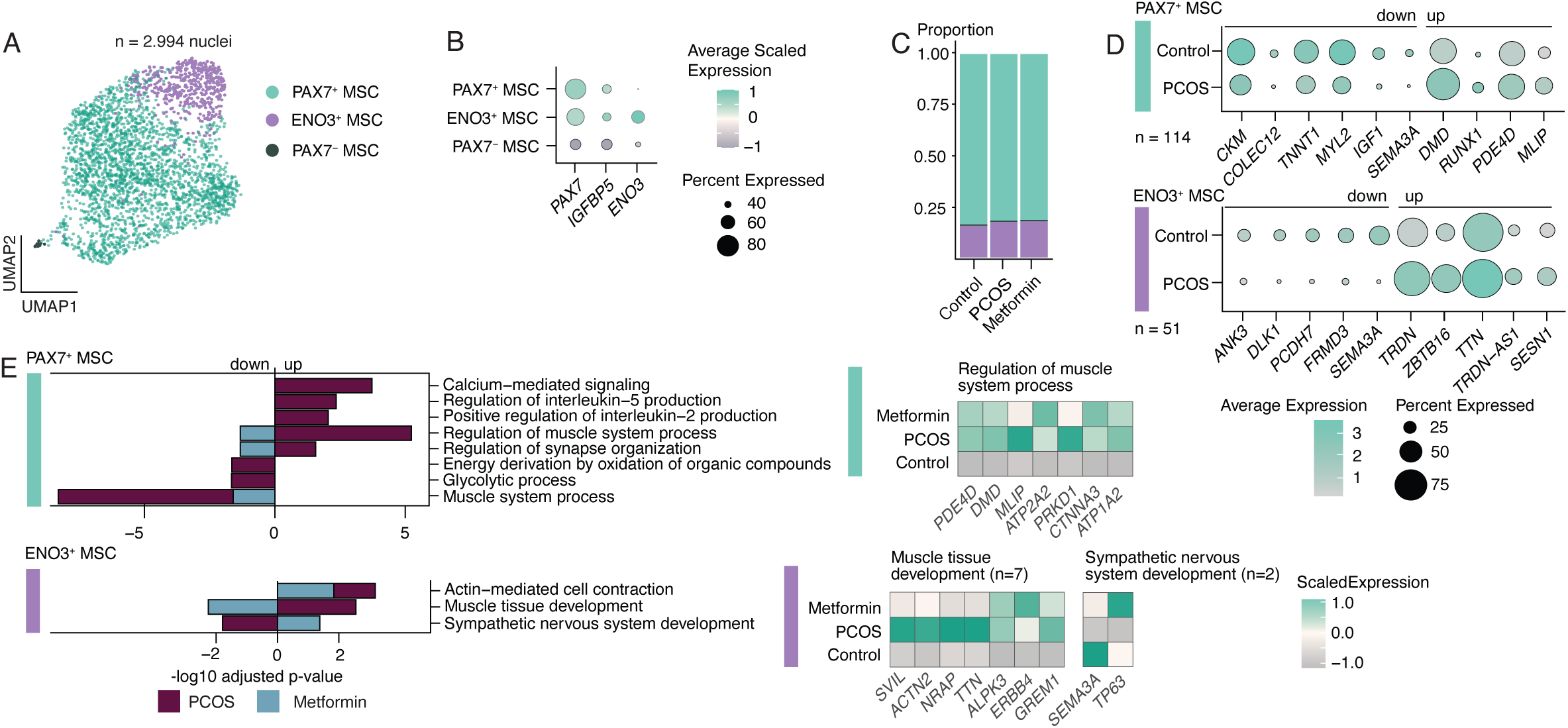
Transcriptomic characterization of satellite cells, and the effect of 16-week metformin treatment in women with PCOS. A) A Uniform Manifold Approximation and Projection (UMAP) of 2,994 nuclei from all patients revealed three distinct satellite cell subclusters: PAX7^+^, ENO3^+^ and PAX7^-^ muscle satellite cells (MSC). B) Dotplot showing log-transformed gene expression of satellite cell marker genes across identified satellite cell types. C) Barplot depicting the proportions of the three-satellite cell subclusters in Control, PCOS and Metformin group. D) Top 5 of up- and downregulated DEGs comparing PCOS vs. control with the total number of DEGs identified in PAX7^+^ and ENO3^+^ subcluster. E) Gene Ontology (GO) enrichment analysis on DEGs within each subcluster at baseline (PCOS) and after 16-week metformin intervention (Metformin). Heatmaps under the GO show the DEGs in corresponding reversed GO terms.

The largest cluster, PAX7^+^ MSC, was the most dysregulated (n = 114 DEGs), followed by ENO3^+^ MSC (n = 51) (Figure 5D, Table S5). We detected no DEGs in PAX7^-^ MSC, likely due to low number of nuclei. Key genes downregulated in PAX7^+^ MSC include *CKM, TNNT1,* and *MYL2*, which play roles in muscle function, as well as *IGF1*, a key growth factor that determined satellite cell proliferation.^63^ Upregulation of *RUNX1, MLIP*, and *DMD* is typically observed in muscles exposed to myopathic damage and activated satellite cells.^64–66^ *SEMA3A* is downregulated in both PAX7^+^ and ENO3^+^ MSC, which may impair slow-twitch fiber specification.^67^

GO term analysis in PAX7^+^ MSC revealed upregulation of regulatory muscle system processes, and synapse organization, both of which were partially reversed by metformin (Figure 5E, Tables S7-S8). Upregulation of calcium-mediated signaling and immune components (regulation of IL5 and IL2), as well as downregulation of energy derivation by oxidation and glycolytic processes were not affected by metformin (Figure 5E). In ENO3^+^ MSC, dysregulation of muscle tissue and sympathetic nervous system development was reversed by metformin. In both PAX7^+^ and ENO3^+^ MSC, dysregulation of muscle system processes and actin-mediated cell contraction, respectively, was further exacerbated by metformin (Figure 5E). Taken together, satellite cells comprise distinct transcriptional subpopulations, with PAX7⁺ cells showing the greatest dysregulation in metabolic and growth signaling, while metformin selectively rescues some pathways but worsens contractile gene disturbances, underscoring subtype-specific effects on satellite cell function.

### Transcriptional signature of immune, endothelial and perivascular subpopulations in PCOS muscle

Skeletal muscle from women with PCOS shows an overall downregulation of immune system related transcripts;^68^ however, data on immune profiling in PCOS skeletal muscle are limited. We characterized 4,714 skeletal muscle immune cells, identifying two macrophage populations (MF and Inflammatory MF [Inf-MF]), B and T cells, as well as mast, and NK cells, based on curated marker genes (Figures S4A-S4B). Immune subpopulation proportions did not differ between women with and without PCOS (Figure S4C). Differential expression analysis was performed across all identified clusters; however, limited number of cells and few DEGs restrict the scope for functional inference. Thus, results are presented descriptively. T cells showed the highest number of DEGs (n = 29), followed by MF (n = 22), mast cells (MAST) (n = 10), Inf-MF (n = 6), and NK cells (n = 5), while no DEGs were detected in B cell subpopulation (Figure S4, Table S5). GO analysis showed pathway enrichment only for T cells, including upregulation in antigen receptor-mediated signaling, lymphocyte proliferation, positive regulation of IL-5 production, and immune response-activating signaling pathways, none of which were affected by metformin treatment (Figure S4E).

In the smaller endothelial and perivascular clusters, we identified 2,013 and 1,712 nuclei respectively (Figures S4F-S4G and Figure S4K-S4L), with no differences in proportions between women with and without PCOS (Figures S4H, S4M). Only capillary endothelial cells (capEC) and pericytes showed dysregulation in PCOS, with 51 and 48 DEGs, respectively (Figures S4K, S4N). CapEC DEGs were enriched in intracellular receptor signaling and long-chain fatty acid transport, with no impact of metformin treatment (Figure S4J).

### Reprogramming of cell-cell communication in PCOS skeletal muscle

Given the enrichment of biological processes related to skeletal muscle function, ECM organization and pro-fibrotic propensity in PCOS skeletal muscle, we next examined whether these transcriptional programs were accompanied by changes in cell-cell communication among subpopulations. The interaction strength, probability, signaling pathways and ligand-receptor pairs are listed in Tables S14-S19.

Differential interaction heatmaps showed the highest number of interactions among endothelial (receivers) and FAP subpopulations (senders), while the strongest differential interaction was between FAPs (senders) and muscle fiber types (receivers) (Figure S5A). Comparison of the global cell-cell communication landscape between control and PCOS skeletal muscle reveals decreased interaction strength in MF-I in PCOS, whereas MF-IIa and MME^+^ FAPs showed increased incoming and outgoing interaction strength (Figures S5B-C). Metformin increased interaction strengths in MF-I and MF-II and decreased MME^+^ FAPs interaction strength (Figures S5B-C), suggesting a reprogramming of communication programs.

Pathway level analysis in different subpopulations revealed that PTPRM, laminin, and collagen pathways exhibited the strongest overall signaling in all three groups, primarily due to signaling in MF-I, MF-IIa and MME^+^ FAPs (Figure 6A). We observed enhanced ANGPTL, SEMA3, WNT, and TGF-β signaling in the FAP subpopulations whereas signaling diminished in MF-I and MF-II. ANGPTL signaling has been linked to metabolic diseases and altered lipid metabolism,^69^ implying that increased ANGPTL signaling could underlie the observed lipotoxicity in PCOS skeletal muscle. Metformin does not reverse the signaling patterns in MF-I and MF-II; however, it does decrease signaling in FAP subpopulations to some extent (Figure 6A).

**Figure 6:**
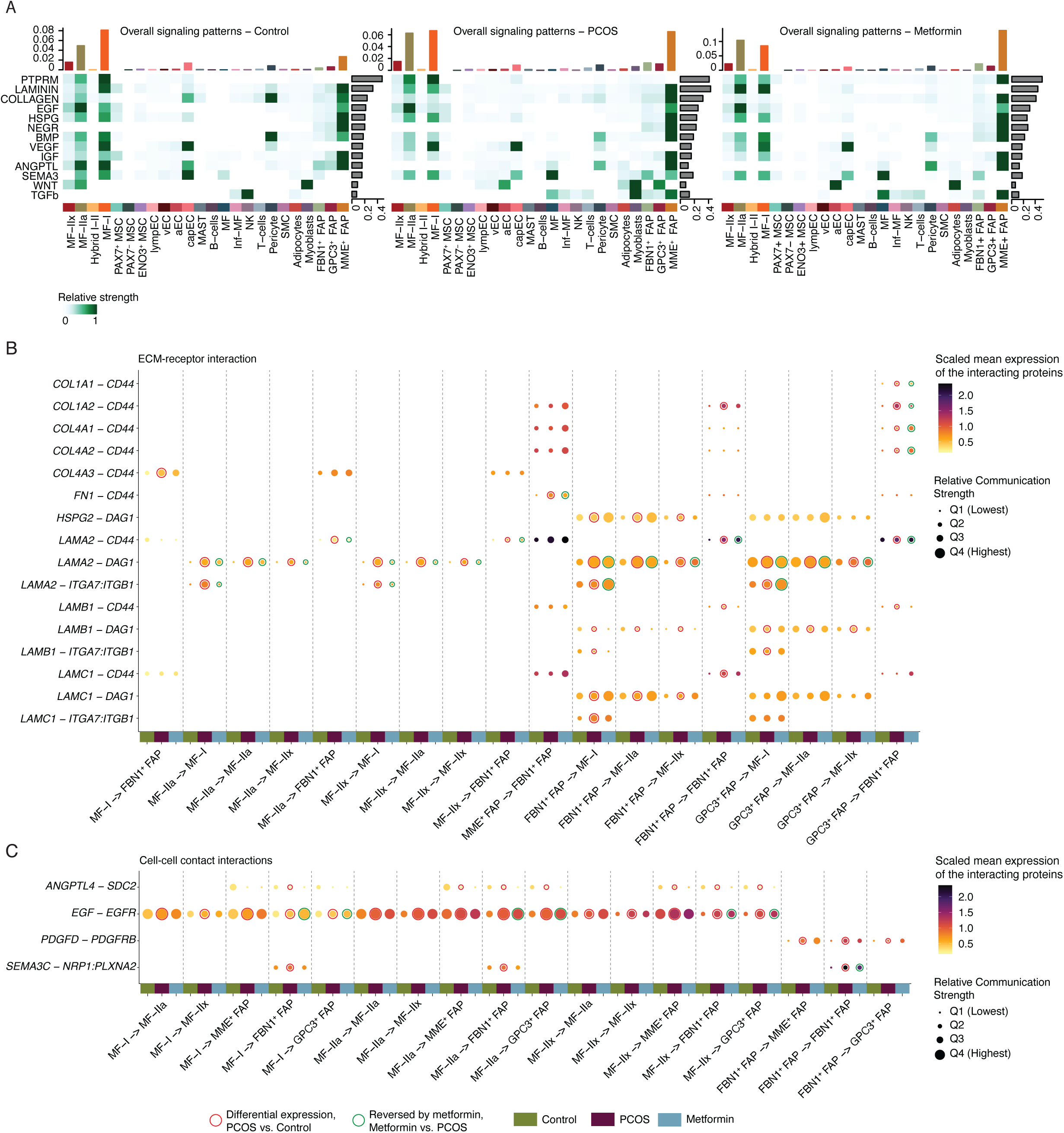
Skeletal muscle – FAPs cross talk and ligand-receptor interaction. A) Heatmaps representing overall signaling patterns across identified pathways in each cell subpopulation in Control, PCOS and Metformin. The bars represent the relative signaling strength of each cell type or pathway, respectively. B) Dotplot of CellChat-predicted extracellular matrix (ECM) receptor interactions across skeletal muscle cell subpopulations in control, PCOS and metformin-treated group. C) Dotplot of CellChat-predicted cell-cell contact interactions. In B and C, the Y-axis shows ligand-receptor pairs and the X-axis shows sending (source) and receiving (target) cell type interaction. Dot color intensity indicates the scaled mean expression of ligand and receptor encoding genes combined. Dot size indicates the relative communication strength binned into quartiles (Q1, lowest, Q4, highest). The colored bar below the X-axis denotes the group. Red circles around the dot indicate interaction where the ligand or receptor is differentially expressed between control and PCOS, or PCOS and metformin. Green circles indicate interaction of which differential expression is reversed by metformin treatment.

To assess whether changes in pathway-level communication were supported by transcriptional alterations, we integrated CellChat and DEG analysis results. Overlapping DEGs as receptor-ligand pairs are presented in Table S17-19. Key findings in PCOS skeletal muscle show dysregulation of ECM-receptor interaction, including cell-to-cell miscommunication between muscle fiber types, with upregulated signaling in MF-IIa, MF-IIx ligand LAMA2 to DAG1 in MF-I, MF-IIa and MF-IIx, and ITGA7:ITGB1 in MF-I, which is reversed by metformin (Figure 6B). Communication from muscle fiber types to FAPs shows upregulated COL4A3 in MF-I to CD44 in FBN1^+^ FAPs, and LAMA2 in MF-IIa and MF-IIx to CD44 in FBN1^+^ FAPs, suggesting that muscle fibers communicate only with FBN1^+^ FAPs. More profound communication was detected from FAPs to skeletal muscle fibers, with HSPG, LAMA2, LAMB1 and LAMC1 in FBN^+^ and GPC3^+^ FAPs to DAG1, ITGA7:ITGB1 on MF-I, MF-IIa, and MF-IIx. Metformin treatment selectively reversed these dysregulations. Upregulated intra-FAP communication through collagens: COL1A1, COL1A2, COL4A1, COL4A2, COL4A3, and to CD44 is reversed by metformin only in the GPC3^+^ to FBN1^+^ direction. Metformin alleviates only FN1–CD44 in the MME^+^ to FBN1^+^ FAPs pair.

Cell-cell contact interaction is upregulated through EGF-EGFR between muscle fiber types and in muscle fiber types to FAPs, with metformin reversal only in FBN^+^ and GPC3^+^ combinations (Figure 6C). Enhanced ANGPTL4 to SDC2 is observed in the direction of muscle fiber types to FAPs communication. FBN1^+^ FAPs communication to all other FAP subpopulation is upregulated with the PDGFD-PDGFRB ligand receptor pair, while intra FBN1^+^ FAPs communication dysregulation is detected through upregulation of SEMA3C-NRP1:PLXNA2, which is reversed by metformin. Together, this interaction network analysis revealed reprogramming of the cell-to-cell communication in PCOS skeletal muscle, with augmented communication between skeletal muscle fibers and FAPs through collagens, laminins and EGF signaling, with selective metformin reversal.

### Skeletal muscle cell-types contributing to disease risk

PCOS shares several genetic loci with type 2 diabetes, fasting insulin (FI), glucose traits, BMI, and waist-to-hip ratio (BMI-adjusted).^70^ We applied CELLEX and CELLECT to our dataset to prioritize cell types contributing to disease risk. Metabolic GWAS for two-hour glucose (2hG), BMI, and WHR showed significant associations with MF-IIx and MF-a, whereas only WHR was linked to adipocytes. Pericytes were prioritized for both T2D and FI, and these were also enriched for MF, Inf-MF, FBN1^+^ and GPC3^+^ FAPs. In the PCOS GWAS a weaker enrichment signal was observed in myoblasts (Figures 7A-B, Table S21).

**Figure 7:**
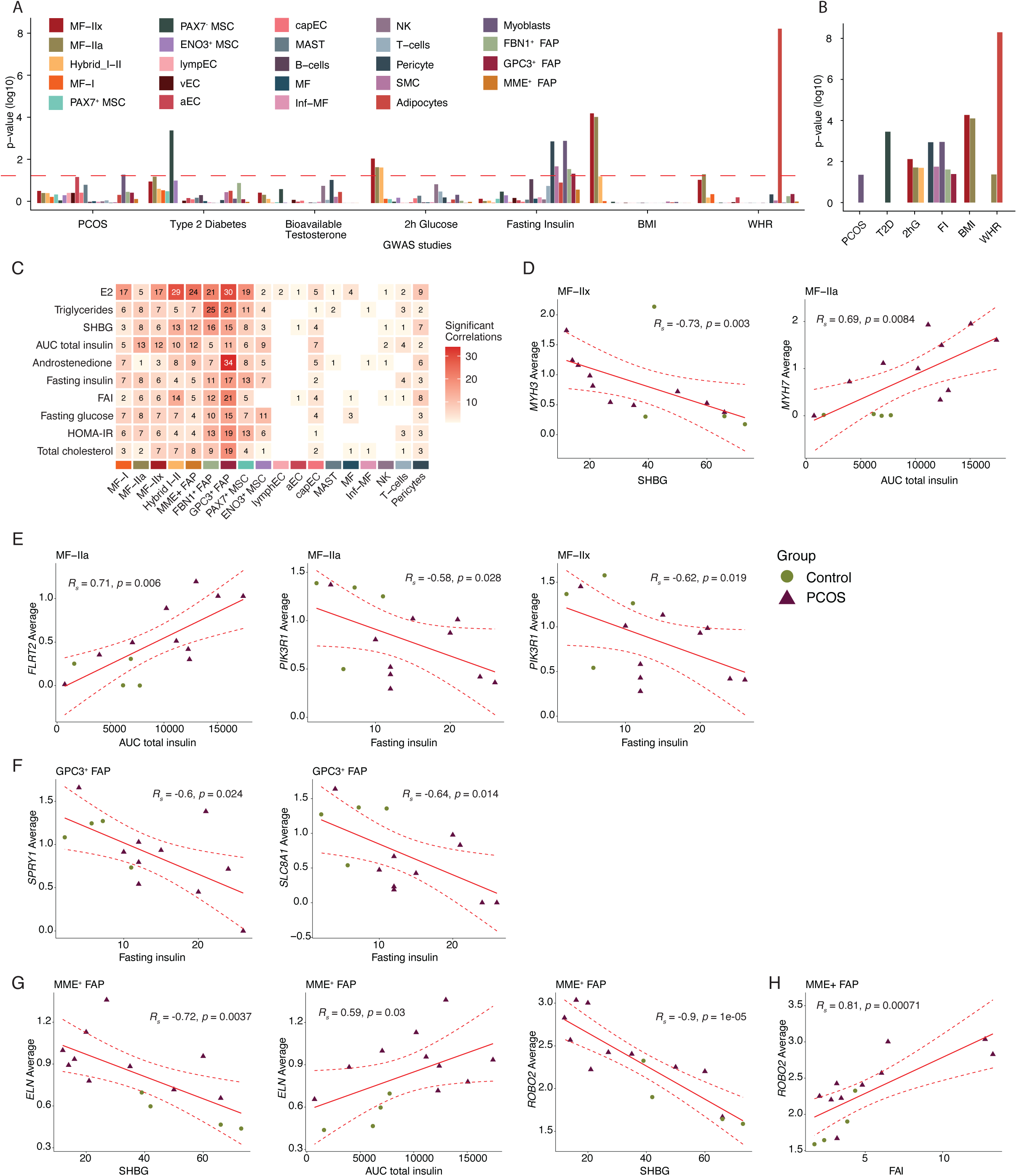
Prioritization of cell types associated with the trait of interest using CELLECT and correlations between DEGs and clinical features. A) CELLECT analysis of the skeletal muscle cell-type enrichment scores derived from GWAS data. The red dashed line indicates significant prioritization after Bonferroni adjusted p-value < 0.05. B) Identified significant subclusters indicate strong associations with genetic variants. C) Heatmap of number of significant Spearman correlations (p < 0,05, Rho > 0.5, Rho < -0.5), between averaged differentially expressed genes in each cell type and selected clinical variables in women with and without PCOS. Number and color intensity indicate the number of significant correlations. D) *MYH3* in MF-IIx negatively correlated with sex hormone-binding globulin (SHBG), and *MYH7* positively correlated with area under the curve (AUC) total insulin. E) *FLRT2* in MF-IIa positively correlated with AUC total insulin and *PIK3R1* in MF-IIa and MF-IIx negatively correlated with fasting insulin. F) *SPRY1* and *SLC8A1* in GPC3^+^ FAP negatively correlated with fasting insulin. G) In MME^+^ FAP, *ELN* negatively correlated with SHBG, and positively correlated with AUC total insulin, and *ROBO2* negatively correlated with SHBG. H) *ROBO2* in MME^+^ FAP positively correlated with free androgen index (FAI). See ‘Statistical analyses for a detailed description of Spearman’s correlations. *RS* = Spearman’s correlation, *p* = p-value.

### Correlation between averaged DEGs per cell type and clinical biochemical traits

To investigate the relationship between cell-type specific alterations and clinical variables that were significantly different between PCOS and controls (Table S1), we performed correlation analysis at baseline across all cell types (Figure 7C, Table S22). In skeletal muscle cells markers of skeletal muscle *MYH3* and *MYH7* negatively correlated with SHBG (R = -0.73, p = 0.002) and positively with area under the curve for insulin (AUCInsulin) (R = 0.69, p = 0.0084), suggesting that both elevated insulin resistance drive changes in muscle phenotype (Figure 7D, Table S22). In MF-IIa, *FLIRT2*, an ECM protein playing a role in cell-cell adhesion,^71^ positively correlated with AUCInsulin (R= 0.71, p = 0.006), while increasing fasting insulin levels negatively correlate with *PIK3R1* in MF-IIa (R = -0.58, p = 0.028) and MF-IIx (R = -0.62, p = 0.019) (Figure 7E), suggesting that women with PCOS might poses an impaired *PIK3R1*, which has been previously linked to severe insulin resistance in patients with SHORT syndrome.^72^ *SLC8A1*, one of the top DEGs, and *SPRY1* in GPC3+ FAPs, negatively correlates with fasting insulin (R = -0.64, p = 0.014, R = -0.64, p = 0.024, respectively) (Figure 7F), linking hyperinsulinemia to dysregulated calcium signaling and disinhibited growth factor pathways. In MME^+^ FAPs, *ELN* expression correlates with both AUCInsulin (R = 0.59, p = 0.03) and SHBG (R = -0.72, p = 0.0037), implying a strong relationship between ECM production and hyperinsulinemia (Figure 7G). *ROBO2* has been previously discovered as key dysregulated gene in PCOS endometrium, which inversely correlated with hyperandrogenism.^73^ Similarly, *ROBO2* inversely correlated with SHBG and positively correlated with FAI in MME^+^ FAPs (Figure 7H), suggesting that hyperandrogenism causes global dysregulations across different tissues. These correlations provide a mechanistic link between the systemic hyperinsulinemic and hyperandrogenic state, and the cell-type specific pro-fibrotic and reprogramming we observe in GPC3^+^ and MME^+^ FAPs.

### Validation of DEGS using antibody-based multiplex tissue profiling

To determine the spatial localization of DEGs and validate if the transcriptomic alterations could be observed at the protein level, antibody-based profiling was performed using an iterative multiplex workflow. A fixed panel of five antibodies outlining different structures in skeletal muscle tissue was developed (Figure 8A). The panel was combined with one antibody at a time targeting a specific DEG, to visualize the expression pattern of specific DEGs in relation to different skeletal muscle cell types (Figures 8B-F). Samples from two controls, two PCOS patients before treatment, and PCOS patients after a 16-week metformin treatment were included. Five DEGs were selected for further analysis using multiplex tissue profiling based on antibody availability and immunohistochemical staining pattern in skeletal muscle in the Human Protein Atlas database (www.proteinatlas.org): COL4A2, COL15A1, ELN, LAMB1 and XIRP2. ELN is positive in fibroblasts, accompanied with weaker staining in both MF-I and M-II fibers (Figure 8B). COL4A2 and COL15A1 mostly overlapped with the fibroblast marker, with COL15A1 showing additional positivity in endothelial cells. For both COL4A2 and COL15A1, the expression appears higher around MF-I fibers compared with M-II fibers (Figures 8C). LAMB1 was distinctly expressed in basement membranes and fibroblasts, with highest expression levels in structures surrounding MF-I fibers especially in PCOS (Figure 8E), which aligns with the CellChat results, where increased LAMB1-DAG1 communication was detected in FAP-myofiber signaling. Finally, XIRP2 showed a variable expression in both MF-I and MF-II fibers, with gradients along the fibers (Figure 8E). Interestingly, there was a tendence towards higher fibroblast expression in the PCOS samples compared to the control, suggesting an upregulation of ELN at the protein level. For the other investigated DEGs, no detectable difference between individuals could be observed at the protein level using antibody-based imaging.

**Figure 8:**
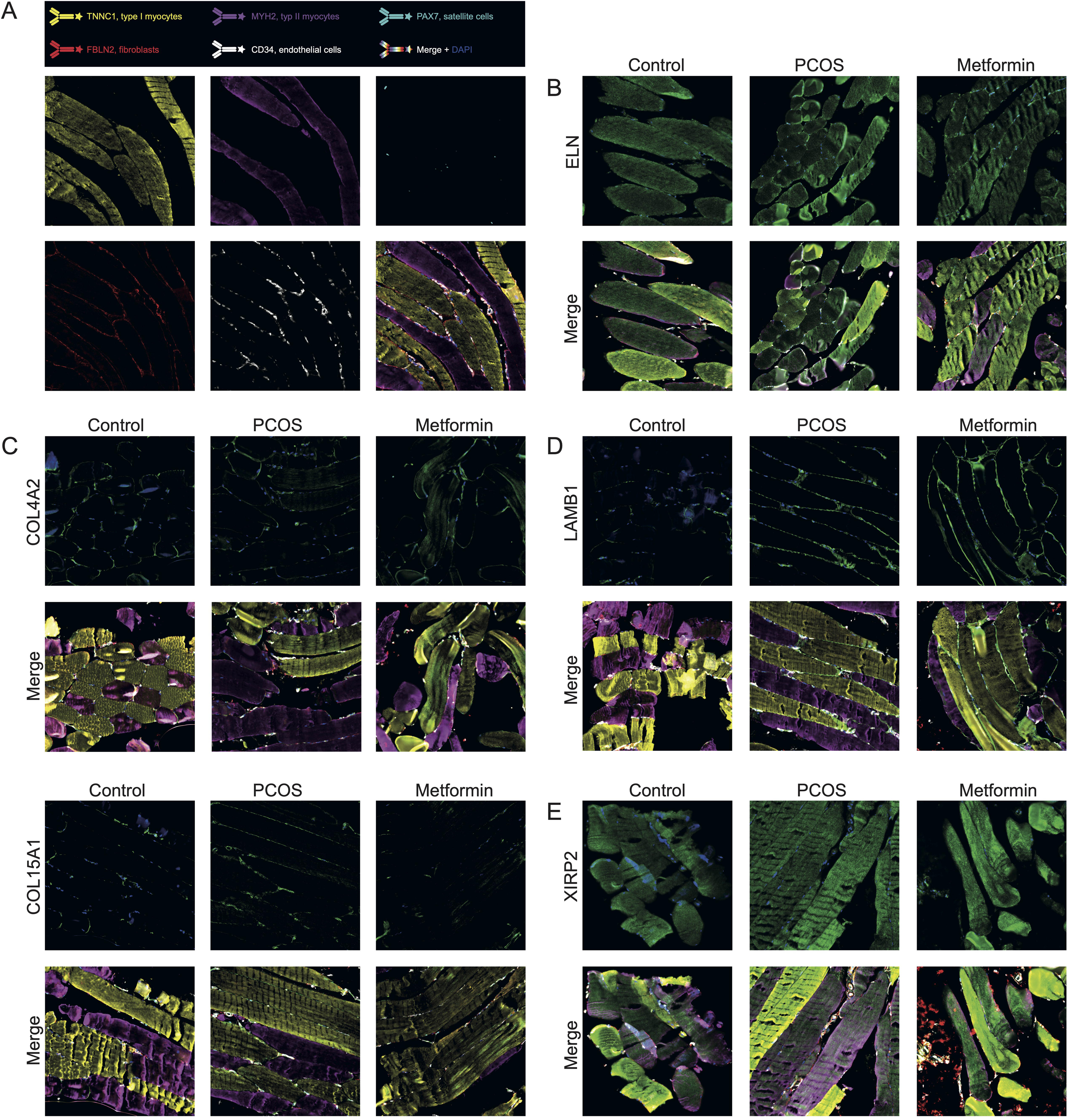
Antibody-based imaging of DEGs using multiplex tissue profiling. A) A fixed 5-plex panel outlines different cell types in skeletal muscle: M-I fibers (TNNC1, yellow), M-II fibers (MYH2, purple), satellite cells (PAX7, cyan), fibroblasts (FBLN2, red), and endothelial cells (CD34, white). B-C) Multiplex staining of five different DEGs is visualized in green (COL4A3, COL15A1, ELN, LAMB1, and XIRP2), with the top panels showing only the DEG together with DAPI, and bottom panels displaying all seven channels (5-plex panel, DEG, and DAPI). Representative images are shown for control samples, PCOS patients before treatment (PCOS week 0), and PCOS patients after 16 weeks of metformin treatment.

## Discussion

PCOS skeletal muscle displays hallmarks of metabolic dysfunction, including insulin resistance, mitochondrial dysfunction, lipotoxicity and dysregulated gene and protein expression.^3,6,7,74^ However, its cellular and molecular heterogeneity and response to metformin remain unclear. In this study, we present the most detailed skeletal muscle atlas to date, analyzing 72,247 nuclei from overweight, hyperandrogenic, insulin resistant women with PCOS, as well as controls with similar age, weight, BMI, and post 16-week metformin intervention. We identify PCOS-specific molecular signatures, dysregulation in metabolic pathways and cell-to-cell communication, and distinct cell-type effect of metformin, providing a mechanistic insight of metformin action in skeletal muscle. This study lays the foundation for functional validation of skeletal muscle cells *in vitro*. These findings reframe our understanding of skeletal muscle dysfunction in PCOS, shifting focus from bulk tissue studies to the cellular interplay between myocytes and FAPs as potential drivers of PCOS metabolic impairment.

### Impaired metabolic flexibility in PCOS skeletal muscle cells

We found that in PCOS, only skeletal muscle subpopulations Hybrid I-II and MF-IIa exhibit mitochondrial dysfunction and increased reliance on fatty acid oxidation, while MF-IIa and MF-IIx display impaired glucose handling, and decreased ROS detoxification, with negligible reversal effect of metformin intervention. The convergence of metabolic dysfunction in fast-twitch fibers aligns with our clinical observations that PCOS is associated with a reduced proportion of oxidative fibers and impaired glycolytic capacity.^75^ The persistence of these metabolic signatures in satellite cell-derived MF-IIx *in vitro* provides evidence that the metabolic and transcriptomic alterations observed *in vivo* persist outside the hormonal milieu of PCOS. These findings are consistent with previous reports that skeletal muscle cells retain disease phenotype *ex vivo*,^76^ suggesting that a degree of PCOS specific metabolic memory is intrinsic (imprinted) and persists independent of circulating androgens. Similarly to single cell data, metformin treatment in MF-IIx myotubes *in vitro* did not reverse mitochondrial dysfunction, nor did we detect any major differences at the transcriptomic level, despite keeping our dose above the therapeutic concentration.^77^ Despite evidence of metabolic memory, PCOS MF-IIx demonstrated similar glucose uptake to controls *in vitro* despite the hyperinsulinemia in PCOS patients, indicating some flexibility in glucose metabolism dysregulation in the absence of the diseased milieu, which has been described previously.^78^ Although we observed dysregulation in both PAX7^+^ and ENO3^+^ satellite cells related to muscle biology and energy metabolism *in vivo*, which was partially reversed by metformin, *in vitro* transcriptomics revealed no differences at Day 0, suggesting that satellite cell alterations may require *in vivo* microenvironment to manifest.

### Pro-fibrotic metabolic remodeling of distinct FAP subpopulations in PCOS

Beyond myofibers, we identify two major FAP subpopulations GPC3^+^ and MME^+^ FAPs, as displaying substantial transcriptional and metabolic alterations, revealing heterogeneity within the stromal compartment that has not previously been addressed in PCOS. Here, we show that while GPC3^+^ and MME^+^ FAPs retain their abundance in PCOS, they undergo a metabolic reprogramming switch to pro-fibrotic phenotype displaying upregulated DEGs and GO terms related to ECM organization and TGF-β signaling. GPC3^+^ FAPs exhibit attenuation of the muscle-supportive phenotype through downregulation of muscle-related genes such as *ARHGEF25* and *MYL2*, fitting into the narrative of a phenotype switch resulting in muscle dysfunction.^40^ Given that MME^+^ FAPs are normally characterized by high adipogenic propensity,^42^ we observed downregulation of adipogenic and immune-related genes such as *CFD*, *TRIM5*, and *CXCL14* alongside altered expression of *SLC8A1* suggesting possible alterations in FAPs metabolic and immunomodulatory capacity. GPC3^+^ FAPs additionally display upregulated BMP signaling, reinforcing the pro-fibrotic FAP phenotype associated with hyperinsulinemic milieu of PCOS.^7,42^ A key finding is that a 16-week metformin intervention exerts selective effects almost exclusively on GPC3+ and MME+ FAPs. Metformin robustly reversed key signaling pathways of fibrosis, while metabolic components like oxidative phosphorylation, Ser/Thr kinase and ERK cascade were not reversed. Our scCellFie analysis further reveals that both GPC3^+^ and MME^+^ FAP subpopulations exhibit a distinctive metabolic signature characterized by enhanced glutamine utilization, coupled with metabolic tasks supporting ECM biosynthesis. Flux analysis has demonstrated that TGF-β increases the incorporation of glutamine-derived carbon into proline, indicating that proline is a major cellular destination for glutamine in myofibroblasts.^79^ Thus, the elevated glutamine metabolic tasks observed in GPC3^+^ and MME^+^ FAPs, along with their upregulated ECM organization programs and suggest that the cells are metabolically equipped to sustain enhanced collagen production. CD90^+^ FAPs, the pro-fibrotic subpopulation enriched in T2D muscle, exhibit elevated glycolytic activity associated with enhanced ECM production,^80^ suggesting that this metabolic reprogramming is common feature of FAP dysfunction across insulin-resistant conditions. Since metformin inhibits TGF-β downstream signaling through AMPK activation,^39^ suppressing fibrotic signaling would reduce collagen production, which we observed as a partial reversal of the upregulated metabolic tasks related to glycosylation and glutamine metabolism.

### Cell–cell communication reveals a pro-fibrotic rewiring in PCOS skeletal muscle

Comprehensive cell–cell communication analysis uncovered a profound rewiring of intercellular signaling in PCOS skeletal muscle, with FAPs emerging as central signaling hubs. MF-I fibers showed reduced interaction strength in PCOS, whereas MF-IIa fibers and MME⁺ FAPs displayed increased signaling activity. Metformin partially reprogrammed this network by restoring interaction strength in MF-I and MF-II fibers and reducing excessive signaling in MME⁺ FAPs, thus potentially ameliorating the dysregulated signaling.

Pathway analysis revealed dominant dysregulation of ECM–receptor interactions, particularly collagen and laminin signaling, primarily driven by MF-I, MF-IIa, and MME⁺ FAPs. Upregulation of collagen and laminin signaling has previously been observed in PCOS endometrium.^73^ Androgen exposure in mice activated ovarian ECM and induced pro-fibrotic remodeling,^81^ which we observed through increased *ANGPTL, SEMA3, WNT,* and *TGF-β* expression in FAPs. This supports an androgen-driven stromal remodeling and a pro-fibrotic phenotype, linked to metabolic dysfunction and lipid handling, which may contribute to altered intercellular signaling and lipotoxicity in PCOS muscle.^8,69^ While metformin had limited impact on restoring fiber-specific pathway signaling, it attenuated aberrant pathway activation within FAPs, suggesting selective targeting of stromal remodeling programs.

Ligand-receptor interaction analysis confirmed transcriptionally supported ECM miscommunication. Muscle fiber-to-fiber miscommunication was characterized by increased LAMA2–DAG1 and LAMA2-ITGA7:ITGB1 signaling, changes that were reversed by metformin. Unlike in muscle dystrophies, where laminin mutations lead to the loss of muscle integrity, ^82^ overexpression and upregulated laminin signaling in PCOS likely lead to ECM stiffness which has been shown to activate pro-fibrotic FAPs state.^83^ Communication from muscle fibers to FAPs was selective, primarily targeting FBN1⁺ FAPs through COL4A3–CD44 and LAMA2–CD44, whereas FAP-to-fiber signaling was markedly amplified, with FBN1⁺ and GPC3⁺ FAPs transmitting HSPG2, laminin, and collagen signals to all fiber types. Thisasymmetry indicates that FAPs act as dominant ECM signal senders, and intra-FAP collagen signaling interactions further support an autocrine pro-fibrotic loop, which is partially reversed by metformin. Together, these findings demonstrate an ECM-centered, FAP-driven pro-fibrotic communication network in PCOS skeletal muscle, with selective but incomplete normalization by metformin.

Correlation analysis between averaged DEGs per cell type and clinical biochemical traits revealed the strongest and most consistent associations in FAPs, particularly the GPC3⁺ and MME⁺ subsets. Notably, *ROBO2*, previously identified as a key dysregulated gene in PCOS endometrium and shown to inversely associate with hyperandrogenism,^73^ exhibited a similar pattern in skeletal muscle MME⁺ FAPs, negatively correlating with SHBG and positively with FAI. These findings support a model in which systemic hyperandrogenism and hyperinsulinemia drive coordinated, cross-tissue transcriptional dysregulation, mechanistically linking endocrine imbalance to the pro-fibrotic remodeling and cellular reprogramming observed in FAP subpopulations.

We detected a small difference in LAMB1 protein levels using multiplex staining, while the other analyzed DEGs appeared similar across samples. Notably, despite a clear fold-change at the RNA level, such differences are not always reflected at the protein level. In addition, antibody-based proteomics has a lower dynamic range compared to mass spectrometry-based methods, and fibroblasts are small structures that are challenging to visualize in a tissue context. Nevertheless, multiplex imaging enabled us to confirm the spatial localization of the DEGs to specific cell types within skeletal muscle tissue. Interestingly, three of the analyzed DEGs (COL4A2, COL15A1, and LAMB1) were more abundant in fibroblasts surrounding MF-I fibers than in those around MF-II fibers, suggesting that fibroblasts in skeletal muscle have distinct properties and regional differences, further emphasizing the need to study alterations at the single-cell type level.

### Limitations of the study

We used snRNA-seq, which captures only nuclear transcripts and may not fully reflect changes in cytoplasmic mRNA. Moreover, snRNA-seq is prone to ambient RNA contamination arising during tissue dissociation. We addressed this using DecontX, however, the complete elimination cannot be guaranteed. Immune cell populations are systematically underrepresented in snRNA-seq compared to single cell RNA-seq, consequently our analysis likely underestimates the contribution of immune cells to PCOS muscle pathology. In this study metabolic dysfunction is inferred from transcript abundance and scCellFie predictions, therefore, further studies are needed to confirm enzyme activity and metabolic flux. Lastly, the cohort included in this study is relatively modest, which may limit generalizability across diverse PCOS phenotypes.

In summary, this is the first cellular atlas of PCOS skeletal muscle, providing a valuable resource for understanding PCOS skeletal muscle dysfunction, deciphering fiber-type-specific metabolic impairment converging with pro-fibrotic FAP reprograming that collectively drive the metabolic sequalae of this syndrome. Uniquely, our study provides the first single-cell resolution map of metformin’s action in human skeletal muscle, uncovering a mosaic of responsive and refractory populations that identifies unmet therapeutic needs and establishes the cellular framework necessary for developing precision therapeutics that target the right cells.

## Resource availability

### Lead contact

Requests for further information and resources should be directed to and fulfilled by the lead contact Elisabet Stener-Victorin.

### Materials availability

The study did not generate any unique reagents.

### Data and code availability

Single-nuclei RNA-seq data have been deposited to KI Data Repository, and are publicly available upon request, as of the date of publication. Accession number is listed in the key resource able.

The code is available at https://github.com/ReproductiveEndocrinologyMetabolism.

## Acknowledgements

We sincerely thank the participants in this study and the research midwives, especially Liselotte Blomberg and Berit Legerstam for the coordination of participants. Computational analyses were enabled by resources provided through the project UPPMAX 2022-003 at Uppsala University.

This work was supported by Swedish Medical Research Council: project no. 2022-00550 (ESV); Distinguished Investigator Grant – Endocrinology and Metabolism

Novo Nordisk Foundation: NNF22OC0072904 and project grant: NNF25OC0104784 (ESV) Diabetes Foundation: DIA2022-708 and DIA2024-860 (ESV); the Regional Agreement on Medical Training and Clinical Research between the Stockholm County Council and the Karolinska Institutet ALFMedN FoUI-973699 and FoUI-1000330 (ESV), Karolinska Institutet Doctoral Education (KID) 2020-00990 and 2023-01283; and the Diabetes Wellness Sverige: IDG-2024-010 (TGS). is supported by a postdoctoral fellowship from SRP Diabetes 2025-00725 (ESV).

## Authors contributions

E.S.-V., Q.D., T.G.S., G.E., C.Li conceived and designed experiments and analysis. A.L.-H. collected skeletal muscle biopsies and screened participants. T.G.S. and C.Li performed sample processing, nuclei isolation and library preparation, and T.G.S. together with G.E. and E.S.-V. analyzed the single nuclei and clinical data. T.G.S, J.R, S.D, H.L. performed satellite cell isolation, differentiation assay, seahorse assay and glucose uptake assay. H.L. performed staining of the myotubes and carried out the imaging. T.G.S. prepared prime seq libraries from RNA isolated by E.L. and analysed the data. C.O. performed sex steroid analysis. R.S. and C.L. performed multiplex tissue profiling. E.S.-V., T.G.S and G.E. wrote the manuscript with contributions from C.Li. All authors read and approved the manuscript.

## Declaration of generative AI and AI-assisted technologies

During the preparation of this manuscript, the authors used AI-assisted tools (Claude.ai) to support editing and language refinement. All AI-generated content was reviewed, edited and verified by authors, who take full responsibility for the integrity and accuracy of the published work. AI tools were not used for data analysis, or generation of figures. The authors declare that no AI tools is listed as an author.

## Supplemental information titles and legends

Document_S1

Figures S1-S5

Tables S1-S22

## Material and methods

### Participants and intervention

Ethical approval has been granted by the Regional Ethical Review Board of Stockholm, Sweden Dnr: 2015/1656-31/2 with amendment Dnr: 2024-00633-02 and is approved by the Medical Products Agency: EUCT: 2024-514505-64-00 and registered at Clinicaltrials.gov: NCT02647827.^84^ The study is conducted in agreement with the Declaration of Helsinki. All women received oral and written information and provided written informed consent prior to participation and were examined at the Women’s Health Research Unit at the Karolinska University Hospital, Stockholm. Participants were between 18 and 40 years of age with a BMI ≥25. PCOS diagnosis was set according to the Rotterdam criteria, excluding any endocrine-related disorders.^85^ Ovarian volume (>10 cm^3^) and number of antral follicles (2-9 mm) was measured with transvaginal ultrasound. In women with PCOS, irregular menstrual cycles were defined as >35 days. Clinical hyperandrogenism was assessed with Ferriman-Gallwey score >4. Control subject had similar age, weight and BMI, regular menstrual cycles (28 days ±2 days), <12 antral follicles (2-9 mm), ovarian volume <10 cm^3^, Ferriman-Gallwey score <4, and no signs of insulin resistance. Exclusion criteria for all women were any medication or hormonal treatment for at least 12 weeks prior to the start, and pregnancy or breastfeeding in the last 6 months.

Sex steroid analyses included dehydroepiandrosterone (DHEA), androstenedione, testosterone, and estradiol measured by gas chromatography-tandem mass spectrometry.^86^ All other hormones were analyzed in an accredited laboratory at Karolinska University Hospital: serum insulin by chemiluminescence using the Elecsys Insulin reagent kit; plasma glucose by enzymatic reaction photometry using the GLU Gluco-quant Glucose/HK kit; sex hormone-binding globulin by electrochemiluminescence using the SHBG kit; triglycerides by enzymatic reaction photometry using the TG Triglycerides GPO-PAP kit; and total cholesterol by enzymatic reaction photometry using the CHOL Cholesterol CHOD-PAP kit. All reagents were from Roche Diagnostics and analyses were performed on the Cobas 8000 (Roche Diagnostics Scandinavia AB).

Biopsies of the vastus lateralis muscle of the quadriceps femoris were collected under local anesthesia using a Weil-Blakesley conchotome (AB Wisex, Mölndal, Sweden) in the morning after an overnight fast on menstrual cycle days 6 - 8. One part of biopsy was used for satellite cell isolation, one part was snap frozen in liquid nitrogen and stored at -80°C, and one piece was fixed in 4% paraformaldehyde for histology.

After the baseline measurements, women with PCOS were randomized to one of three interventions for 16 weeks using our online electronic case report form: (1) lifestyle management alone; (2) metformin + lifestyle; or (3) electroacupuncture + lifestyle. In brief, lifestyle management for all women with PCOS included an initial meeting with information about the importance of healthy eating, regular physical activity and weight management. In addition, they received a book with lifestyle advice and received a weekly text message reporting their steps over the previous week and whether they had menstrual bleeding. The metformin group received 500 mg orally, 3 times per day for 16 weeks. The dose was increased from 500mg per day the first week, to 1,000mg per day the second week, to the full dose 1,500mg per day, the third week. For details see published protocol.^84^ snRNA-seq and *in vitro* experiments were performed on skeletal muscle biopsies collected at baseline and after 16 weeks of intervention in the (2) metformin + lifestyle group.

### Nuclei extraction of frozen human skeletal muscle biopsies

Nuclei were extracted from 20-30g of frozen skeletal muscle tissue using Chromium Nuclei Isolation Kit with RNase Inhibitor (10x Genomics) according to the protocol. In brief, the biopsies were lysed in Lysis Buffer and dissociated with plastic pestille until homogenous, followed by nuclei collection in the Nuclei Isolation Column at 16000rcf for 20 seconds. Debris was removed by resuspension of nuclei in Debris Removal Buffer followed by centrifugation at 700rcf for 10 minutes and washing steps using Wash and Resuspension buffer. At the end nuclei were concentrated in 30µl of Wash and Resuspension buffer, counted and visualized to ensure the nuclei are well preserved.

### Single-nuclei RNA-seq library preparation and sequencing

Library construction was performed using Chromium Next GEM Single Cell 3’ Gel Bead Kit (Dual Index) v3.1 (10X Genomics) and Chromium Next GEM Chip G Single Cell to capture 10,000 nuclei per reaction. Library quality was verified with 2100 Bioanalyzer (Agilent) and sequencing was performed on the Illumina NovaSeq6000 system at Novogene, UK, with the objective of achieving a minimum coverage of 30,000 raw reads/nucleus (Table S3).

### Single-nuclei RNA-seq analysis

The raw data from the FASTQ files were processed and aligned to a reference transcriptome (GRCh38-2020-A) including introns using Cell Ranger Software (v.6.1.1). Filtered reads from Cell Ranger were used for the quality control and each sample was converted into a Seurat (v4) object before filtering.^87^ Nuclei with < 300 genes and genes expressed in less than 3 nuclei were removed as well as those with > 5% mitochondrial reads, > 5% ribosomal reads and > 1% hemoglobin reads and duplicates were removed with scDblFinder. We implemented SingleCellExperiment framework, applied DecontX that estimates the contamination proportion, and adjusts the counts accordingly.^88^ Then, SelectIntegrationFeatures with 2000 features and FindIntegrationAnchors with default parameters were used to identify the anchors while IntegrateData was used for the integration and sctransform (v2.0) was used for the normalization and clustering. Principle component analysis (PCA) and uniform manifold approximation and projection (UMAP) were used for dimension reduction, and the count matrix was normalized and transformed using Seurat Log-Normalize.

#### Cell type annotation and subclustering

Cell type annotation was performed manually based on the extensive literature curation and applying FindAllMarkers to the sctransformed clusters which resulted in the most highly expressed genes in each cluster. The six main clusters skeletal muscle cells, FAPs, satellite cells, immune cells, perivascular cells, and endothelial cells were further subsetted, re-integrated as described before and annotated manually as in the previous step.

#### Differential gene analysis and Gene Ontology (GO) enrichment analysis

FindMarkers and the MAST method were used to identify DEGs in the contrasts: Control vs. PCOS, PCOS vs. metformin group, using the Bonferroni-adjusted P value < 0.05, log_2_FC > 0.5 and genes detected in > 10% of the nuclei in both populations.^89^ Based on log_2_FC, the genes were categorized into either upregulated or downregulated before subjected to the functional enrichment analysis for biological processes ontology using clusterProfiler.^90^ The most enriched biological processes with a false discovery rate (FDR) < 0.05 were reported and the most relevant terms were visualized, whereas in the contrast PCOS vs. metformin only matched (reversed) GO terms were selected.

#### Cell-to-cell communication analysis with CellChat

To computationally predict and analyse cell-to-cell communication networks, CellChat (v2.1.2) was applied on the Seurat object labelled with all subpopulations.^91^ Cellchat objects of Control, PCOS and Metformin grouped sampled were generated and processed seperatelly with default CellChat setting, where communication probabilities were calculated with computeComunProb with population.size to consider subpopulation proportions. Cell-to-cell communications were predicted based on ligand and receptor expression calculated by gene expression per subpopulation, where information flow between ligand and receptors was labelled as outgoing signaling indicative of ligand expression, or incoming signaling being indicative of receptor expression. The CellChat objects were then merged into two groups; Control versus PCOS and PCOS versus Metformin, and the interactions were compared using the compareInteractions to identify interaction with differing relative interaction strengths between groups. Interactions of differentially expressed ligand and receptors were identified by filtering the data using the MAST-based DEG data generated previously between the groups, allowing for the identification of ligand-receptor pair with impaired gene expression. Interactions and their associated pathways with high relative strength and ligand-receptors pairs with differentially expressed genes, were selected for further analysis.

#### Metabolic tasks inferred from snRNA-seq

To characterize cell-type-specific metabolic programs in PCOS skeletal muscle, we employed scCellFie (v.0.5.0)^35^ integrated with Scanpy (v.1.11.5)^92^ in Python (v.3.10) to infer metabolic task activity based on gene-protein-reaction rules. Gene expression was smoothed using KNN averaging k = 10, and α = 0.33. To compute metabolic gene scores, we implemented precomputed thresholds as previously described.^35^ Prior to differential metabolic analysis (DMA) we verified balanced representation of cells across experimental groups in each cell subpopulation, to avoid bias in effect size estimation. DMA was performed within each cell type using Wilcoxon rank-sum tests with Benjamini-Hochberg correction. Metabolic tasks were considered significant at FDR < 0.05 and Cohen’s d > 0.2.

#### Candidate etiologic cell types

To investigate which cell types are more relevant to specific diseases or traits, we examined genetic signal from nine GWAS studies (Table S20) and linked it to the cell types based on the estimated gene expression specificity using CELLECT (v1.3).^93^ First, gene expression matrix was converted into gene expression specificity estimates using CELLEX (v1.2.2). Then, the matrix was used together with GWAS summary statistics as an input for CELLECT-MAGMA covariate analysis using the default parameters. Significant cell types were identified using Bonferroni P value threshold < 0.05.

#### Correlation analysis

The relationship between averaged gene expression of DEGs per subpopulations with clinical variables that differed between women with and without PCOS per sample, was investigated with Spearman’s rank correlation between the samples, of which relationship with *R_S_* > 0.5 or *R_S_* < -0.5 and *P* < 0.05 were deemed significant.

### Human satellite cell isolation and differentiation

To validate snRNA-seq results and confirm that the skeletal muscle cells maintain the features PCOS *in vitro*, we isolated and differentiated human satellite cells into myotubes (n_PCOS_= 6, n_Ctrl_= 6).^94^ One day after the sample collection, skeletal muscle tissue was minced and enzymatically digested (Ham’s F10 medium (Gibco), Collagenase type IV (Sigma-Aldrich), trypsin and 0.5% BSA) at 37°C with regular shaking for the total of 10-15 minutes. Growth media (GM; Ham’s F10 (Gibco), 20% FBS (Gibco), 1% PEST) was added to stop the digestion, then the solution was filtered through a 40µm strainer and centrifuged at 800g for 7 minutes. The pellet was resuspended in 5ml of GM and plated on a small petri dish. To ensure the separation of the fibroblasts, the cells were incubated in the petri dish for 3 hours in the incubator (37°C, 5% CO_2_) and then transferred to a new petri dish. The GM was changed after 4 days, then every second day.

Myogenic differentiation was performed to confirm the differentiation capacity of the isolated cells. Upon expansion, the satellite cells were seeded in 24-well plate and chamber slides (Corning) with density of 0.1 x10^6^ cells/well and when sub-confluent treated with intermediate media (IM; DMEM 1g/L glucose (Gibco), 10%FBS (Gibco), 1%PEST) when sub-confluent, followed by differentiation media by differentiation media (DM; DMEM 4.5g/L glucose (Gibco), 2% horse serum (Gibco), 1% PEST) for 6 days. Upon differentiation, the supernatant was collected on the last day and stored at -20°C for further experiments. On day 0 and day 8 the cells were washed with PBS and treated with 700µl QIAzol (Qiagen) for RNA extraction.

#### Prime-seq – library preparation

Upon RNA isolation from myotubes with ReliaPrep™ RNA Miniprep (Promega), 10 µg of RNA/sample was used for the library construction following Janjic et al.^95^ In brief, RNA, RT mix (Maxima H Minus RT, Maxima RT Buffer 5X (ThermoFisher), dNTP (NEB), TSO (E3V7NEXT, sigma), ddH_2_O) and Barcoded oligodT (E3V7NEXT, Sigma-Aldrich) were mixed and incubated at 42°C for 1,5h, followed by pooling of the samples and a cleaning step with clean-up beads in 22% PEG (PEG 8000 1.1g (Sigma-Aldrich), NaCl (5M) 1ml, Tris-HCL (1M, pH 8.0), Igepal – CA630 (10% solution 5µl, Sigma-Aldrich), Sodium Azide (10% solution 25µl), H_2_0) at a 1:1 ratio of the reaction to beads. The tube was placed on a magnet and washed with 1ml 80% ethanol twice. The beads were dried and incubated in 17µl ddH_2_O followed by elution. In the digestion step, 2µl of 10X Exol Buffer (NEB) and 1µl of Exonuclease I (NEB) was added to the cDNA followed by incubation (20 minutes at 37C, 10 minutes at 80°C), cleaning in clean-up beads at a ratio of 1:0.8 and concentrating in 20 µl of ddH_2_O. Upon the elution, we performed pre-amplification (KAPA HiFi 2X RM (Kapa Biosystems), PreAmp primer SINGV6 (IDT), ddH_2_O), with the following PCR setup: initial denaturation at 98°C for 3 minutes; 10 cycles of denaturation (98°C for 15 seconds), annealing (65°C for 30 seconds), elongation (72°C for 4 minutes); and a final elongation at 72°C for 10 minutes. The DNA was cleaned with clean-up beads at 1:0.8 ratio, concentrated in 10µl and quantified using Qubit 1X dsDNA HS Assay Kit (Thermo Fisher, #Q33A31)). Fragmentation and ligation were performed on 7.8ng/µl using NEBNext Ultra II FS Library Preparation Kit (NEB) according to the manufacturer’s instructions. The library size was selected using SPRI-select beads (Beckman Coulter) at 1:0.5 and 1:0.2 ratio, followed by amplification with Q5 Master Mix (NEB), 1µl Index primer i5 (5µM, TruSeq i5), 1µl Index primer i7 (5µM, Nextera i7) using the following setup: 98°C for 30 seconds: 11 cycles of 98°C for 10 seconds, 65°C for 75 seconds; and 65°C for 65 minutes. Double size selection was performed once more as before, followed by a quantification using 2100 Bioanalyzer (Agilent) with High Sensitivity DNA Analysis kit (Agilent).

#### Sequencing and data analysis

Sequencing was performed on Illumina NovaSeq Plus Series (NovoGene) at 36G raw data per sample. The quality check was performed using fastQC, followed by trimming and mapping using zUMIs pipeline, which uses STAR to perform alignment to the human genome (GRCh38 v48).^96–98^ After gene alignment, UMIs were deduplicated and the count matrix was generated. Generated gene counts were used to perform downstream analysis in R (v4.3.2).^99^ Only gene counts > 10 were included in the analysis. DESeq2 (v1.42.1) was used for differential expression analysis using the Wald test.^100^ Genes exhibiting statistically significant alterations in expression (adjusted p value < 0.1, L_2_FC > 0.5), were identified as DEGs. PCA analysis was performed on variance stabilizing transformation (vst). Biological Processes (BP) Gene Ontology was performed using the clusterProfiler (v4.6.2) on overlapping and unique DEGs and GO terms were selected (adjusted p value < 0.05) and plotted.^90^

### Myotube oxygen consumption

To examine the impact of PCOS on skeletal muscle metabolism, we measured the oxygen consumption rate (OCR) using Seahorse XFe96 Analyzer. Satellite cells were seeded (1 x 10^4^ cells/well) and differentiated into myotubes in the Seahorse XFe96 cell culture microplate and were treated with IM ± 1 mM metformin (Fisher Scientific) for 2 days and with DM ± 1mM metformin for 6 days. On the day of the assay, cells were washed with the Seahorse XF assay media (glucose (10mM), sodium pyruvate (1mM), L-glutamine (2mM)) (Agilent Technologies) and equilibrated for an hour at 0% O_2_. OCR calculations were performed under basal conditions and after the sequential addition of oligomycin 1µM, FCCP 1µM, and rotenone/antimycin A 1µM/1µM. After the experiment, the myotubes were washed with PBS and lysed using RIPA lysis buffer to quantify total protein content for normalization using the Qubit protein assay (Invitrogen) according to manufacturer’s instructions.

#### Glucose uptake

To determine how PCOS and metformin treatment influence glucose uptake we performed Glucose Uptake-Glo^TM^ Assay (Promega). 1 x 10^4^ satellite cells/well were seeded in the white opaque 96 well-plates and after 24h the GM was changed to IM ± 1 mM metformin for 2 days and with DM ± 1 mM metformin for 4 days. The day prior to the detection, DM was changed to DM without serum and metformin. Next day the cells were incubated in no glucose DMEM (Gibco) ± 1µM insulin (Fisher Scientific) and without metformin for 1h at 37°C. The media was removed and 50µl of 1mM 2DG (2-Deoxyglucose) in PBS was added and incubated at 25°C for 45 min. 25µl Stop Buffer, 25µl Neutralization Buffer, and 100µl of 2DG6P Detection Reagent were added, followed by incubation for 1h at 25°C and luminescence recording on a SpectraMax® iD3s luminometer (Molecular Devices). Data were normalized based on the number of cells seeded.

#### Immunofluorescence

Differentiated myotubes in Nunc Lab-Tek II chamber slide system (Thermo Scientific) were washed with PBS and dried prior to 10-minute fixation in 4% paraformaldehyde. Slides were washed in washing buffer (PBS, 1% Tween 20 (Sigma Aldrich)) and permeabilized in 0.5% Triton X-100 (Sigma Aldrich) for 5 minutes following by another washing step. Blocking buffer (10% goat serum in PBS) was applied for 20 minutes followed by primary antibody cocktail treatment overnight (BA-F8-s (1:100, DSHB), SC-71-s (1:250, DSHB), 6H1-s (1:100, DSHB)). The following day, slides were washed and incubated with the secondary antibody cocktail (Goat anti Mouse IgG2b Secondary Antibody, Alexa Fluor 594, Goat anti-Mouse IgG1 Secondary Antibody, Alexa Fluor 647, Goat anti Mouse IgM Heavy Chain Secondary Antibody, Alexa Fluor 488 (1:500, Invitrogen) for 1 hour. Nuclei were co-stained with Hoechst (1:1000) and the slides were mounted with Vectrashield® Vibrance™ (Bionordica).

### Multiplex tissue profiling

#### Panel development

A fixed 5-plex panel was built to target five different structures in skeletal muscle tissue: MF-I fibers, MF-II fibers, satellite cells, fibroblasts and endothelial cells. Protein markers and corresponding antibodies were selected based on (i) literature search, (ii) previously tested reliable antibodies with available data in the Human Protein Atlas (www.proteinatlas.org), and (iii) immunohistochemical staining pattern in test slides of human skeletal muscle. Suitable antibodies were first tested with a single-plex run, i.e. one antibody at a time, to ensure compatibility with the OPAL detection system, and only continuing with antibodies that generated a similar staining pattern as observed by regular immunohistochemistry. Then, each antibody was tested in each position of the multiplex workflow to find the best location for each antibody to ensure a stable panel, taking into consideration the following parameters: (i) antibody specificity to the intended structure, (ii) signal-to-noise ratio, and (iii) signal strength. Finally, when the best position for each antibody had been selected, all five panel markers were stained simultaneously in a 5-plex staining, to confirm non-overlapping signals. Primary antibodies for the panel were: anti-PAX7 (AB_528428, Developmental Studies Hybridoma Bank, Iowa City, IA), anti-CD34 (M7165, Agilent Technologies Inc., Santa Clara, CA), anti-FBLN2 (HPA001934, Atlas Antibodies AB, Bromma, Solna, Sweden), anti-TNNC1 (HPA044848, Atlas Antibodies AB), and anti-MYH2 (M4276, Sigma-Aldrich, Saint Louis, MO).

#### Slide pretreatment

Validation of the candidate DEGs was performed on the FFPE skeletal muscle blocks from the single nuclei cohort including samples from 2 Controls, 2 PCOS and 2 Metformin. The blocks were cut in 4µm sections, mounted on adhesive slides and baked at 60°C for 1 hour. Slides were deparaffinized in Xylene, followed by hydration in graded alcohols and blocking for endogenous peroxidase in 0.3% hydrogen peroxide. Antigen retrieval was performed by boiling in pH 6 buffer (Agilent Technologies Inc.) at 125 °C for 4 min using a Biocare Medical Decloaking Chamber PLUS (DC2008INTL). After cooling to 90 °C, the slides were rinsed in deionized water (1−2 min) and transferred to TBS-Tween wash buffer (Thermo Fisher Scientific, TA-999-TT, Waltham, MA, USA) until further processing. To reduce autofluorescence, slides were incubated in a bleaching buffer containing 1.5% hydrogen peroxide, 0.2 M glycine, and 1× TBS-Tween (Thermo Fisher Scientific, TA-999-TT) in 50 mL Falcon tubes. Tubes were rotated under an overhead LED light at RT on a bench for 1 h using a Stuart SRT9D roller mixer. Slides were subsequently washed and stored in TBS-Tween until further use.

#### Multiplex staining

Candidate DEGs proteins were stained one at a time together with the fixed 5-plex panel as follows: Tissue sections were blocked with UV Blocking Buffer (10 min) (UltraVision LP HRP kit, Epredia, Kalamazoo, MI), incubated with primary antibody (30 min), rinsed in TBSTween, incubated with HRP-conjugated secondary antibody (preconjugated ready-to-use) (10 min) (UltraVision LP HRP kit, Epredia), and washed again in TBS-Tween. Slides were then incubated with the cycle-specific OPAL fluorophore (Akoya Biosciences, Marlborough, MA) for 10 min, rinsed, and subjected to HIER (90 °C, pH 6 buffer, (Agilent Technologies Inc.) 20 min). The order of the Opal dyes and panel markers from first to last cycle was: OPAL690 for PAX7 (satellite cells), OPAL620 for CD34 (endothelial cells), OPAL520 for candidate DEG protein, OPAL570 for FBLN2 (fibroblasts), OPAL480 for TNNC1 (MF-I fibers), and OPAL-DIG-780 for MYH2 (M-II fibers). Primary antibodies for the DEGs were: COL4A2 (C1926, Sigma-Aldrich), COL15A1 (HPA017913, Atlas Antibodies AB), ELN (HPA018111, Atlas Antibodies AB), LAMB1 (sc-17763, Santa Cruz Biotechnology, Dallas, TX), and XIRP2 (HPA074599, Atlas Antibodies AB). After each cycle, slides were rinsed and stored in wash buffer. The final step involved incubating slides with OPAL 780 fluorophore-conjugated anti-DIG antibody (1:125 in Epredia Lab Vision Antibody Diluent OP Quanto, #TA-125-ADQ) for 1 h, followed by staining with DAPI (1:1000, Invitrogen, D1306, Thermo Fisher Scientific) for 5 min. Slides were rinsed, mounted with Invitrogen ProLong Glass Antifade Mountant, covered, and cured overnight. Finally, slides were digitized at 40× magnification using PhenoImager HT (Akoya Biosciences).

### Statistical analyses

Statistical analysis on clinical variables was performed in IBM SPSS Statistics (30.0.0.0.), using Mann Whitney U test in the baseline comparison and Wilcoxon signed rank test to investigate changes from baseline to after 16-week metformin intervention in women with PCOS. To detect differences in nuclei proportions in each major cluster, we used Kruskal-Wallis with Bonferroni correction. Statistical analysis for myotube oxygen consumption and glucose uptake data was performed on GraphPad Prism (v. 10.6.1) using one-way ANOVA with Tukey correction and Mann-Whitney test, respectively. Statistical methods used for computational analysis of single nuclei RNA-seq data are described in the corresponding sections above.

