## Supplementary figures and images for "Cell type–resolved transcriptomic map of skeletal muscle in women with polycystic ovary syndrome"

### Supplemental Figure 1

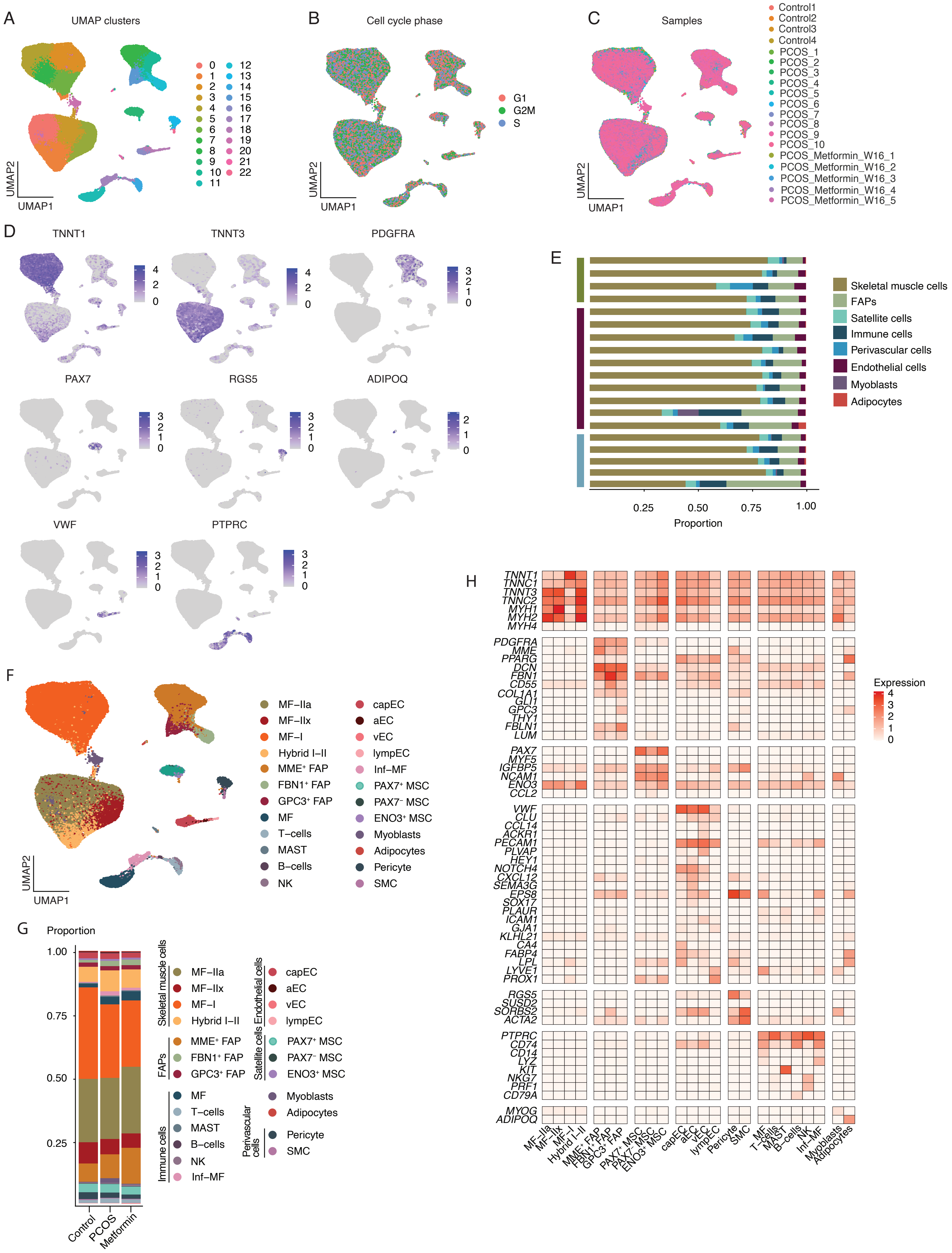

### Supplemental Figure 2

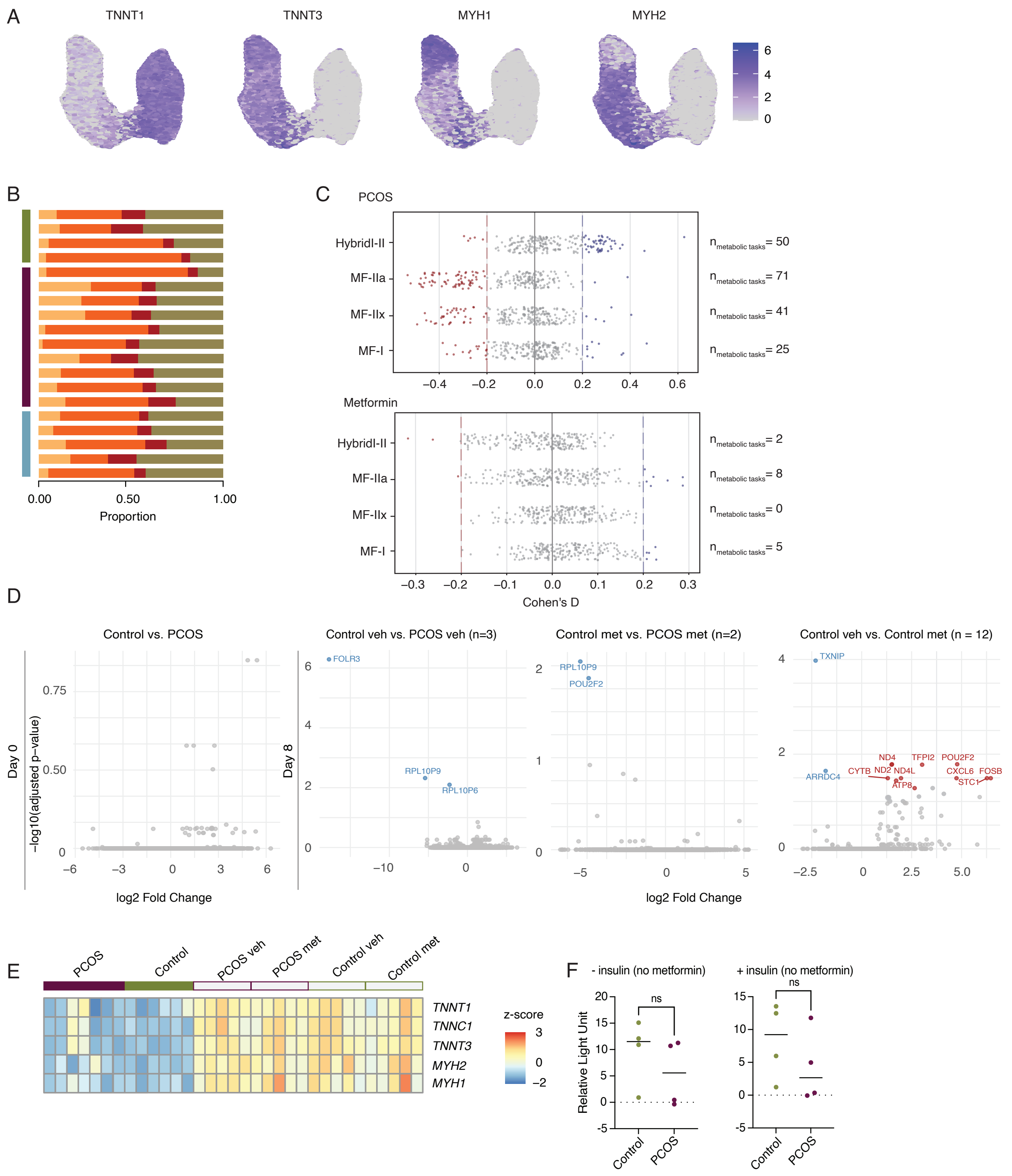

### Supplemental Figure 3

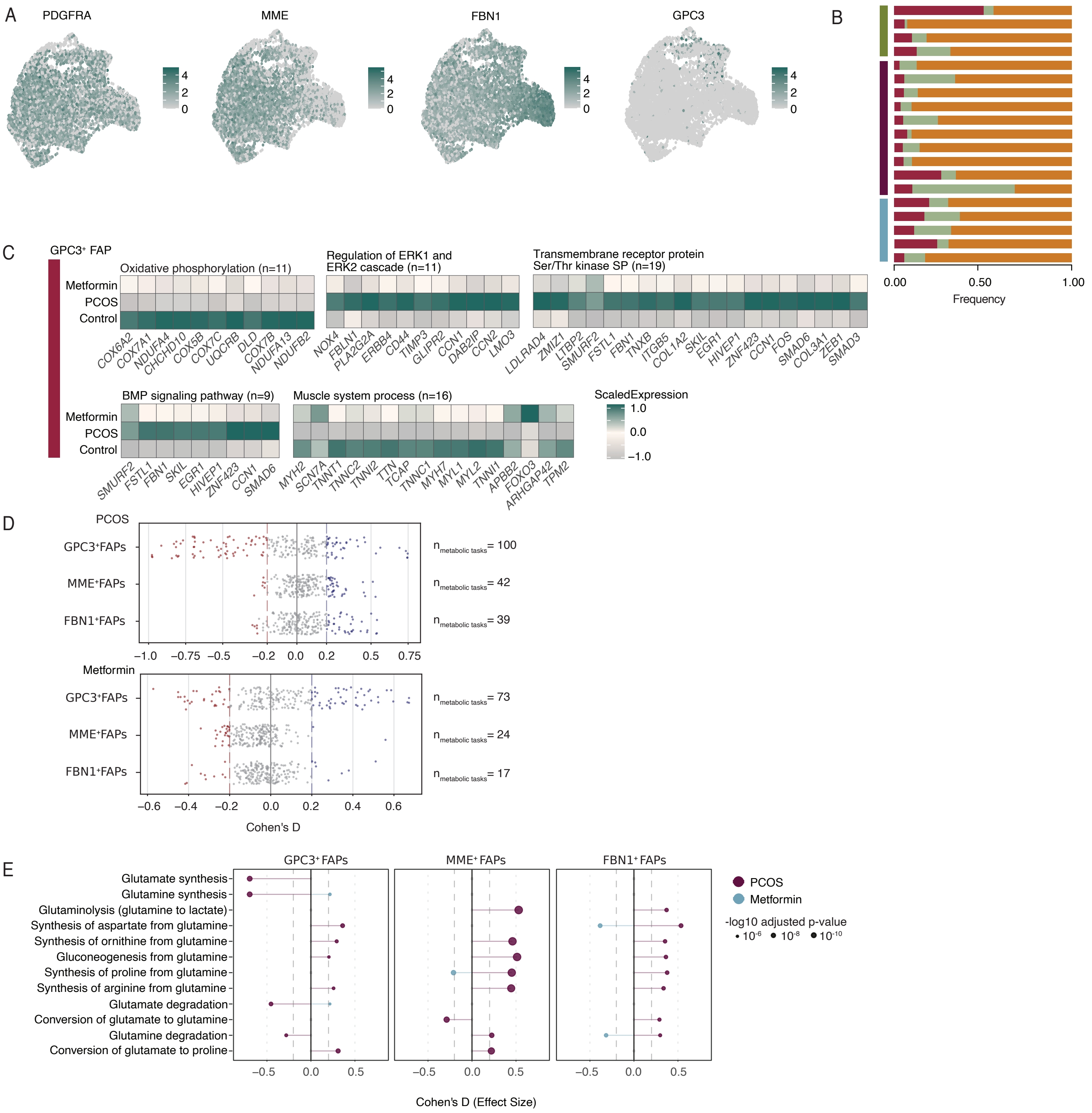

### Supplemental Figure 4

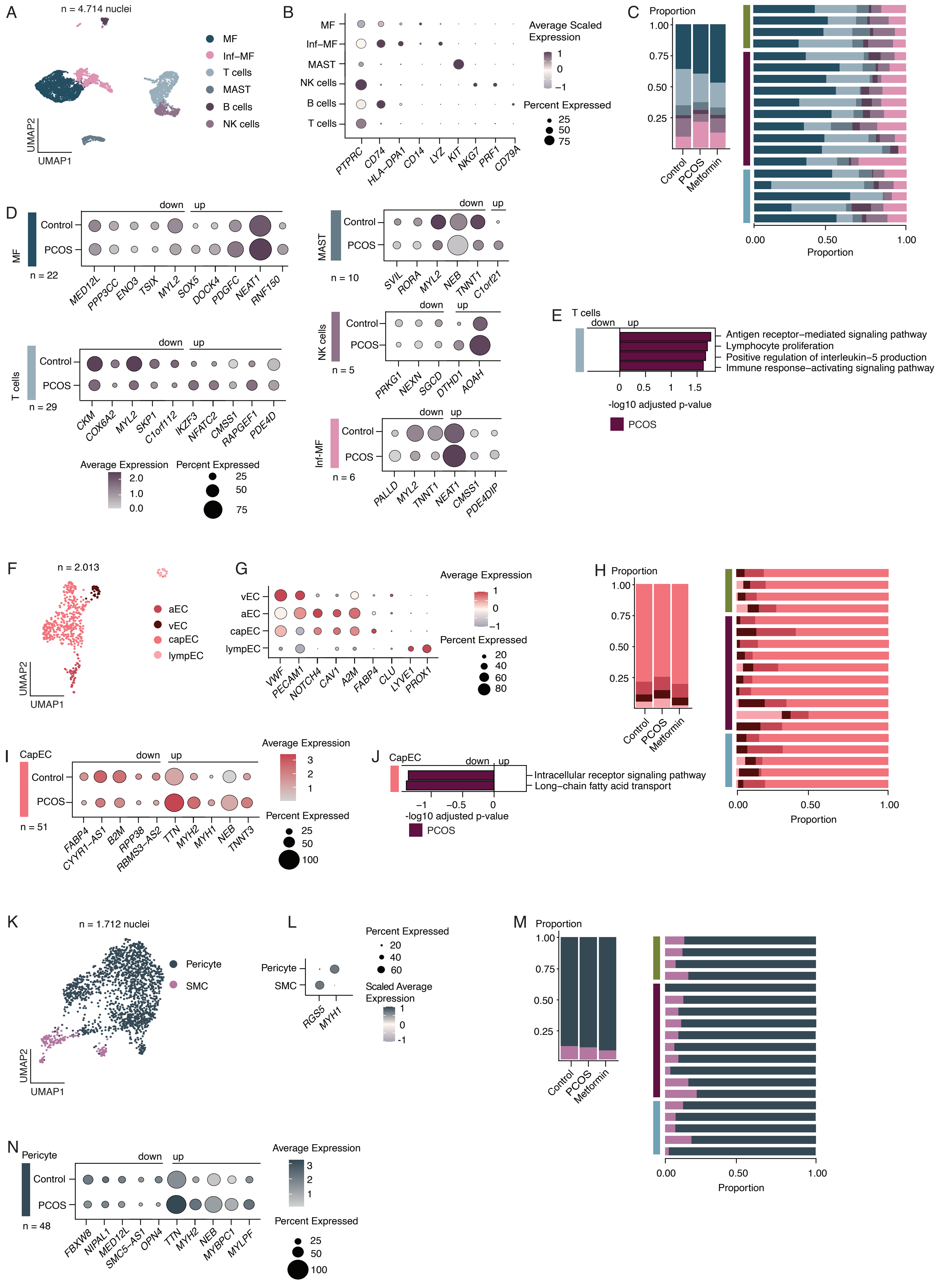

### Supplemental Figure 5

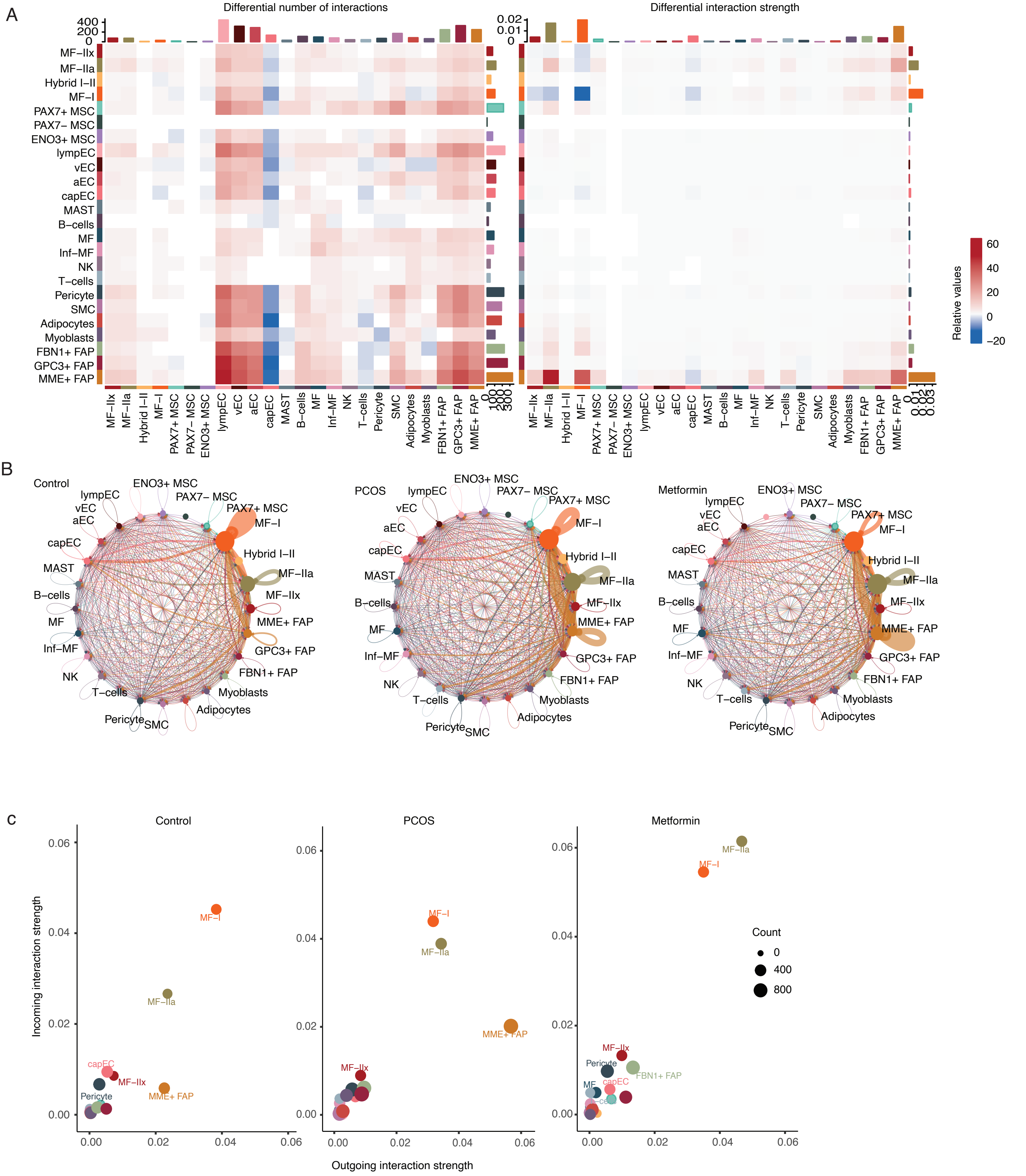
