## Supplemental Figure Legend for "Cell type–resolved transcriptomic map of skeletal muscle in women with polycystic ovary syndrome"

**Figure S1: Single cell profiling of individual samples and quality control**. A) Uniform manifold approximation and projection (UMAP) of integrated snRNA-seq data of 72,247 nuclei showing UMAP clusters, B) cell cycle phase and C) individual sample IDs. D) Feature plots identifying major cell clusters based on the marker genes. E) Bar plot depicting the proportions of each major cell cluster in each sample. Green – Controls, purple – PCOS, blue – Metformin. F) UMAP with projected identified skeletal muscle cell subpopulations and G) their proportions in each group. H) Heatmap of all gene markers used to identify cellular subpopulations. Color indicates average expression.

**Figure S2: snRNA-seq quality control and scCellFie results for skeletal muscle cluster and *in-vitro* validation on 2D myotubes**. A) Feature plots identifying skeletal muscle fibers based on the marker genes. B) Barplot depicting distribution of proportions of each muscle type fiber in individual sample. C) Beeswarm plots visualizing differential metabolic tasks from scCellFie analysis in PCOS (PCOS vs. Control comparison) and Metformin (Metformin vs. PCOS comparison). Each dot represents a significant metabolic task (adjusted p-value < 0.05), positioned based on the Cohen’s d effect size (x-axis). Positive values indicate upregulation; negative values indicate downregulation. Colored points indicate tasks meeting significance threshold (Cohen’s d > ± 0.2), their number is listed at the right side of the plot. D) Vulcano plots depicting significant DEGs from the prime-seq analysis at Day 0 and Day 8 in the corresponding comparisons and the number of DEGs. E) Heatmap showing expression of muscle markers in the satellite cells Day 0 and myotubes at Day 8 with and without metformin. F) Glucose uptake measured in Relative Light Unit in Control (n=5) and PCOS myotubes (n=5), with and without insulin. Ns – not significant.

**Figure S3: extended FAPs snRNA-seq characterization**. A) Feature plots identifying FAPs subpopulations based on marker gene expression. B) Barplot depicting distribution of FAPs subpopulations in individual sample. C) Heatmaps of DEGs from the GO terms, that were not reversed by metformin. D) Beewswarm plots visualizing differential metabolic tasks from scCellFie analysis in PCOS (PCOS vs. Control comparison) and Metformin (Metformin vs. PCOS comparison). Each dot represents a significant metabolic task (adjusted p-value < 0.05), positioned based on the Cohen’s d effect size (x-axis). Positive values indicate upregulation; negative values indicate downregulation. Colored points indicate tasks meeting significance threshold (Cohen’s d > ± 0.2), their number is listed at the right side of the plot. E) Lollipop plot showing metabolic tasks activity differences related to glutamine metabolism across FAPs subpopulations between PCOS (PCOS vs. Control) and Metformin (Metformin vs Control). X-axis is Cohen’s d effect size, size of the dots represents statistical significance (-log10 adjusted p-value).

**Figure S4: Transcriptomic characterization of immune, endothelial and perivascular clusters and the effect of 16-week metformin treatment in women with PCOS**. A) UMAP of 4,714 nuclei from all patients revealed 6 immune subclusters: macrophages (MF), inflammatory macrophages (Inf-MF), T-cells, MAST cells, B-cells, and NK cells. B) Dotplot showing log-transformed gene expression of satellite cell marker genes across identified immune cell types. C) Barplot depicting the proportions of immune cell subclusters in Control, PCOS and Metformin group (left) and individual samples (right). D) Top 5 of up- and downregulated DEGs (PCOS vs. Control) with the total number of DEGs identified in immune cell subclusters. E) Gene Ontology (GO) enrichment analysis on DEGs within T-cell subcluster at baseline (PCOS vs. Control). No reversed GO terms were detected in Metformin group. F) UMAP of 2,013 nuclei from all patients revealed four endothelial cell subclusters: capillary endothelial cells (capEC), arterial EC (aEC), venous EC (vEC), and lymphatic (lympEC). G) Dotplot showing log-transformed gene expression of endothelial cell markers genes. H) Barplot depicting the proportions of the endothelial cell subclusters in Control, PCOS and Metformin group (left) and individual samples (right). I) UMAP of 1,712 nuclei from all patients revealed two perivascular cell subclusters: pericytes and smooth muscle cells (SMC). J) Dotplot showing log-transformed gene expression of perivascular cell markers genes. K) Barplot depicting the proportions of the perivascular cell subclusters in Control, PCOS and Metformin group (left) and individual samples (right). L) Top 5 of up- and downregulated DEGs (PCOS vs. Control) with the total number of DEGs identified in capEC and pericytes. M) Gene Ontology (GO) enrichment analysis on DEGs within each subcluster at baseline (PCOS vs. Control). No reversed GO terms were detected in Metformin group.

**Figure S5: Cell-chat cell to cell interactions**. A) Heatmaps showing differential number of interactions and their strength among the cell subpopulations. The top colored bars represent the sum of incoming signal, whereas tight colored bars represent the sum of outgoing signal. B) Circle plot showing the interaction strength across the cell subpopulations in Control, PCOS and Metformin. C) Scatter plot with outgoing (x-axis) and incoming (y-axis) interaction strength identifying differential patterns in cellular communication between Control PCOS and Metformin.
